# Efficient in vivo genome editing mediated by stem cells-derived extracellular vesicles carrying designer nucleases

**DOI:** 10.1101/2021.02.25.432823

**Authors:** Sylwia Bobis-Wozowicz, Karolina Kania, Kinga Nit, Natalia Blazowska, Katarzyna Kmiotek-Wasylewska, Milena Paw, Elzbieta Karnas, Agnieszka Szyposzynska, Malgorzata Tyszka-Czochara, Olga Woznicka, Dariusz Boruczkowski, Claudio Mussolino, Paweł P. Łabaj, Axel Schambach, Zbigniew Madeja, Toni Cathomen, Ewa K. Zuba-Surma

**Author notes:** Correspondence and requests for materials should be addressed to S.B.-W. or E.Z.-S.

## Abstract

Precise genome editing using designer nucleases (DNs), such as zinc finger nucleases (ZFNs), transcription activator-like effector nucleases (TALENs) and the clustered regularly interspaced short palindromic repeat/Cas9 (CRISPR/Cas9) system, has become a method of choice in a variety of biological and biomedical applications in recent years. Notably, efficacy of these systems is currently under scrutiny in about 50 clinical trials. Although high DNs activity in various cell types *in vitro* has already been achieved, efficient *in vivo* genome editing remains a challenge. To solve this problem, we employed stem cells-derived extracellular vesicles (EVs) as carriers of DNs. We used umbilical cord-derived mesenchymal stem cells (UC-MSCs) and induced pluripotent stem cells (iPSCs) as EV-producer cells, since they are both applied in regenerative medicine. In our proof-of-concept studies, we achieved up to 50% of EGFP marker gene knockout *in vivo* using EVs carrying either ZFN, TALEN or the CRISPR/Cas9 system, particularly in the liver. Importantly, we obtained almost 50% of modified alleles in the liver of the experimental animals, when targeting the *Pcsk9* gene, whose overexpression is implicated in hypercholesterolemia. Taken together, our data provide strong evidence that stem cells-derived EVs constitute a robust tool in delivering DNs *in vivo*, which may be harnessed to clinical practice in the future.

## Introduction

Precise genome editing has become an indispensable procedure in basic research, biotechnology and, more recently, in therapeutic applications. The genome editing toolbox comprises of customizable, site-specific designer nucleases (DNs), such as zinc finger nucleases (ZFNs) [1], transcription activator-like effector nucleases (TALENs) [2,3] and the most recently discovered, clustered regularly interspaced short palindromic repeat/Cas9 (CRISPR/Cas9) system [4,5]. These tools act as molecular scissors and induce DNA double-strand breaks at a predefined site in the genomic DNA. The resulting free DNA ends are then rejoined by the cellular DNA-repair machinery, either by precise homologous recombination or by error-prone non-homologous end joining, to restore genome integrity [6]. Depending on the availability of sequences homologous to the targeted site, the activity of DNs may result in the occurrence of insertion or deletion (indel) mutations at the target locus or precise editing. The latter can be explored for example to correct a disease-associated mutation. Thus, DNs constitute useful tools for genome modification, which can be exploited in the modelling of human diseases, new drug development and can ultimately be harnessed for treatment of genetic diseases [7].

Despite their high activity and precision, effective delivery of programmable nucleases to certain target cells and particularly direct *in vivo* delivery, remains a challenge and hampers their clinical use. Currently, the most effective way of *in vivo* gene targeting is achieved by using viral vectors, which, as natural infective agents, easily transmit foreign genetic material to cells. However, the use of viral components in medical applications is burdened with serious risks, including their immunogenicity and/or malignant transformation of the transduced cells [8,9]. Other available non-viral systems, such as liposomes or other artificial vesicles, usually fall short in effectivity. Although recently developed advanced lipid formulations showed high rate of *in vivo* genome editing [10,11], the use of lipid particles has certain limitations. Such systems often activate the immune response in the host, which leads to their clearance from an organism and ultimately impedes a desired biological effect [12]. Therefore, to overcome these obstacles, we have developed a novel delivery system of hybrid nucleases to cells, based on the use of extracellular vesicles (EVs) derived from stem cells.

EVs are natural nano-vesicles released by virtually any type of cells upon activation and in steady-state. They are composed of a cell membrane and bioactive cytoplasmic components, such as small RNAs, proteins and lipids. EVs play an important role in cell-to-cell communication by transferring their bioactive content to other cells [13,14]. Based on their size and origin EVs are typically divided into: i) small vesicles (exosomes) with a diameter of 30-100 nm, which are derived from endosomal compartment and are characterized by the presence of tetraspanins such as CD9, CD63, CD80; ii) large vesicles (ectosomes; microvesicles; shedding vesicles) with a diameter of 100-1000 nm, which are generated by budding of cell plasma membrane and carry on their surface markers typical for their parental cells; iii) apoptotic bodies, with a diameter over 1000 nm, which are released from a cell during programmed cell death and contain nuclear components, including fragments of genomic DNA [15].

Utility of EVs in various biomedical applications has gained considerable interest in recent years [16]. We have previously shown that EVs derived from stem cells, including umbilical cord-derived Mesenchymal Stem Cells (UC-MSC-EVs) and induced Pluripotent Stem Cells (iPSC-EVs), are capable of transmitting their cargo to target cells, thus affecting their fate and behavior [17-19]. Due to their acellular nature and lack of nucleus, EVs are considered safer than their cellular counterparts, which constitutes a great advantage for future clinical use [19,20].

In this work, we employed EVs derived from human stem cells as delivery vesicles for DNs. We used two human stem cell types: UC-MSCs and iPSCs, both of which are considered for regenerative therapies worldwide. In our proof-of-concept system based on disruption of *EGFP* gene encoding for enhanced green fluorescent protein, we were able to show high editing efficiencies *in vivo*, reaching almost 50% in the liver of the experimental animals. Importantly, when targeting mouse proprotein convertase subtilisin/kexin type 9 (*Pcsk9*) gene, whose overexpression disturbs lipid homeostasis in the liver and leads to hypercholesterolemia [21], we obtained approximately 50% of modified alleles in the animal livers.

Concluding, in this study we proved that stem cells-derived EVs constitute an attractive technological solution to efficiently deliver DNs *in vivo*. Because our system is devoid of any viral component and utilizes EVs from clinically relevant cell types, it can be considered safe and can serve as a basis for future applications in gene therapies for patients affected with genetic diseases.

## Materials and Methods

### Cell culture

UC-MSCs were isolated from umbilical cords obtained from the Polish Stem Cell Bank (Warsaw, Poland) using an explant method described previously [17] and were maintained according to the approval of local ethical committee. Cells were cultured in DMEM/F12 (Gibco/Thermo Fisher Scientific, Waltham, MA, USA) supplemented with 10% fetal bovine serum (FBS; Sigma-Aldrich/Merck, St. Louis, MO, USA) and penicillin (100 U/ml), streptomycin (100μg/ml) solution (P/S; Gibco).

HEK293T cell line was purchased from ATCC and was cultured in DMEM (high glucose) (Sigma-Aldrich), supplemented with 10% FBS (Gibco) and P/S solution (Gibco).

Both cell types were re-plated upon reaching 70% confluency onto new cell culture dishes using TrypLE Select Enzyme solution (Gibco).

hiPSCs were purchased from Gibco (Human episomal iPSC line obtained via reprogramming of CD34+ hematopoietic stem cells with Sendai viral vectors expressing factors: OCT4, SOX2, KLF4, c-MYC). Cells were cultured in Essential 8 medium (Gibco) supplemented with P/S (Gibco) on recombinant human vitronectin (50μg/ml; Gibco) coated plates. Every four days cells were re-seeded on new plates using 0.5 mM EDTA (Invitrogen/Thermo Fisher Scientific) and supplemented with 10 µM/ml Rho-associated protein kinase inhibitor (Y-27632, Merck) for the first day.

All cell types were grown in standard conditions at 37 °C, 5% CO_2_ in a humidified cell incubator.

### Plasmids and genetic modification of cells

Expression plasmids encoding single domains of ZFN, TALEN or CRISPR/Cas9 targeting EGFP gene were obtained by subcloning of a nuclease domain from original plasmids (pRK5.nG(17-2).Neeai_0998, pRK5.nG(18-2).hNqkiv_0999 – for ZFN-L/R; pVAX_CMV_TALEN_GFP_L5_WT-Fok_1459, pVAX_CMV_TALEN_GFP_R5_WT-Fok_1460 – for TALEN-L/R and pU6-GFP-gRNA, pCMV-Cas9-PGKpuro – for gRNA and Cas9) into pLV-EF1a-Nuclease-PGKpuro, or pLV-U6-gRNA-PGKpuro. Stable UC-MSC-DN cell lines expressing a single nuclease domain were generated by nucleofection of 1 µg of an endotoxin-free expression vector into 10^5^ UC-MSCs using the Neon Transfection system (Invitrogen) in the Neon Tip10 with the following parameters of a single electric pulse: 1400V, 30 ms. Immediately after nucleofection, the cells were transferred to pre-warmed cell culture medium supplemented with 10% FBS, without antibiotics. To obtain stable cell lines, the cells were cultured in the presence of 0.8 µg/mL of puromycin for 6 days and expanded afterwards.

HEK293T cells and UC-MSC expressing EGFP gene were generated by transduction with EGFP expressing lentiviral vectors (LV). Viral particles were produced in HEK293T packaging cell line by triple transfection with psPAX2 (Addgene #12260; http://n2t.net/addgene:12260; RRID:Addgene_12260) and pMD2G (Addgene #12259; http://n2t.net/addgene:12259; RRID:Addgene_12259) packaging plasmids and expression vector EGIP (Addgene #26777; http://n2t.net/addgene:26777; RRID:Addgene_26777) [22] using Lipofectamine2000 (Invitrogen) as a transfection agent. The cells were transduced with LV particles at low multiplicity of infection (MOI: 1-2) in the presence of 10 µg/mL of polybrene (Merck). HEK293T-EGFP were selected with 1 µg/mL of puromycin, expanded as single cell-derived clones and genotyped. UC-MSC-EGFP^dim^ cells were sorted out using FACS Aria III cell sorter (BD Biosciences).

To target mouse *Pcsk9* gene two gRNAs were tested. gRNA sequences were *in vitro* synthesized using the GeneArt Precision gRNA Synthesis Kit (Invitrogen). 300 ng of gRNA was mixed with 3 µg of the True Cut Cas9 Protein (Invitrogen) in 12 µL of buffer R from the Neon Transfection System and incubated for 10 min at room temperature. The RNP complexes were introduced to mouse embryonic fibroblasts (MEFs) by a single pulse nucleofection at 1400V, 30 ms using the Neon Tip10 in triplicates. The cells were immediately placed into a pre-warmed DMEM high glucose (Sigma) supplemented with 15% FBS and without antibiotics. Two days after nucleofection the cells were harvested for analysis. The CRISPR/Cas9 lentiviral plasmids targeting mouse *Psck9* gene were obtained by oligo cloning of guide RNA (gRNA) sequences into the CMV-hspCas9-T2A-puro-H1-gRNA vector (Systems Biosciences). The obtained plasmids were used for LV production, as described above. hiPSCs were transduced with LV particles at MOI of 5 in the presence of 5 µg/mL of polybrene (Merck). Stably transduced cells were obtained upon selection with 0.3 µg/mL of puromycin for 6 days. Oligonucleotide sequences used for Pcsk9-gRNA synthesis and cloning to LVs are listed in the Supplementary Table 6. All plasmids are available upon request.

### Genotyping

Genomic DNA from genetically modified UC-MSCs, HEK293T and hiPSCs and control cells was extracted with the GeneMATRIX gDNA Isolation Kit (Eurx) and was used for PCR-based detection of the integrated transgenes. All reactions were performed with Phire Hot Start II DNA polymerase kit (Thermo Scientific). Primers used in the reactions are listed in the Supplementary Table 1.

Copy number of EGFP gene in HEK-EGFP single cell-derived clones was measured with the TaqMan CopyNumber Reference Assay, human, RNase P and calculated with the CopyCaller Software v2.0 (both from Applied Biosystems). K562-EGFP cell line with a single integration of EGFP expression cassette [23] was used as a normalizer.

### Immunofluorescence

hiPSCs cultured on a glass bottom 24-well plate (Eppendorf) were washed with PBS and fixed with 4% paraformaldehyde (PFA; Sigma-Aldrich) for 15 min (RT), followed by permeabilization with 0.2% Triton X-100 solution (Sigma-Aldrich) for 10 min (RT). Fixed samples were incubated with blocking solution composed of 3% BSA (Sigma-Aldrich) in PBS for 1 h at RT, and immunolabeled with anti-human antibodies for PE-conjugated OCT4, FITC-conjugated CD133, or FITC-conjugated SSEA4 (all from BD Biosciences), overnight at 4°C. Cells were further washed twice with PBS. Next, cell nuclei were stained with DAPI (4’,6-diamidino-2-phenylindole, dilactate; 2μM, Life Technologies). Cells were visualized in PBS with calcium and magnesium (Gibco) using Leica DMI6000B inverted microscope equipped with a dry 10x, NA 0.25 objective, digital DFC360FX CCD camera (both from Leica Microsystems) and standard DAPI/GFP/TRITC fluorescence filter combination. Image acquisition and processing were performed with Leica Application Suite X software.

### Isolation of extracellular vesicles

UC-MSC-DNs were washed twice with PBS to remove FBS and cells were kept in DMEM/F12 supplemented with 0.5% bovine serum albumin (BSA; Sigma-Aldrich) and P/S for 48 h to collect EVs. hiPSCs were cultured in Essential8 medium and conditioned media were collected when cell density reached approximately 70-90%. EVs were isolated according to sequential centrifugation protocol, as previously described [17]. Briefly, CMs were centrifuged at 2000 g for 15 min at 4°C, then subjected to ultracentrifugation at 100 000 g, in Sorwall WXUltra 80 Ultracentrifuge equipped with a T-865 fixed angle rotor (Thermo Scientific), for 70 min, 4°C twice, with an intermediate washing step in PBS. EV pellets obtained from approximately 90 mL CM were re-suspended in 150-200 µL PBS (Lonza). Protein concentration was determined with the Bradford assay.

### Nanoparticle Tracking Analysis

EVs were analyzed and quantitated by the NS300 nanoparticle analyzer with the NTA software (NanoSight, Malvern, Worchestershire, UK). Samples were diluted 1:1000 in PBS for optimal particle count. Data were collected with the camera level of 12 and the detection threshold of 3. Three 60-second videos were recorded for each sample using the script control function.

### Transmission Electron Microscopy

Negative staining of EVs was achieved by adsorption of 20 μl of EVs suspended in PBS to a nickel-coated grids (Agar Scientific, Stansted, UK) for 30 minutes, then fixing for 5 minutes with 2,5% glutaradehyde solution. After removing excess liquid with filter paper the samples were stained with 2% uranyl acetate for 30 minutes, washed three times in distilled water for 1 minute and dried. Finally, EVs were visualized by JEOL JEM2100 HT CRYO LaB6 transmission electron microscope (JEOL, Peabody, MA, USA).

### Flow cytometry

Expression of *EGFP* gene was analyzed in cells using BD LSRFortessa and FACS Diva software (BD Biosciences). High resolution cytometry was applied to analyze surface markers expression on EVs with the A50-Micro Flow Cytometer (Apogee Flow Systems). Briefly, EVs suspended in PBS (Lonza) were stained with the SYTO® RNASelect™ Green Fluorescent cell Stain solution (Molecular Probes/ThermoFisher Scientific) and selected antibodies (APC-labeled mouse anti-human: CD45, CD90, CD105, CD309 and CD147, all from BioLegend) for 30 min in the dark. The percentage of positive events was calculated using Apogee Histogram software (Apogee Flow Systems).

### Western blot

Cell pellets and EVs solutions (in 3:1 ratio) were lysed in RIPA buffer (Sigma-Aldrich) containing the Halt Protease Inhibitor Cocktail (Thermo Scientific). 15 µg of protein extracts were separated by Mini-PROTEAN TGXPrecast Gels (BioRad, Hercules, CA, USA) and transferred to PVDF membranes using Trans-Blot Turbo RTA Mini PVDF Transfer Kit (BioRad). Proteins were detected with rabbit polyclonal HA epitope Taq (NB600-363, Novus Biologicals), mouse monoclonal anti-Cas9 (7A9-3A3, Cell Signaling Technology), mouse monoclonal exosome anti-CD9 (10626D, Invitrogen), mouse monoclonal anti-CD81 (MA513548, Invitrogen), goat polyclonal anti-Flotillin-1 (PA518053, Invitrogen), goat polyclonal anti-Syntenin/SDCBP (PA5-18595, Invitrogen), goat polyclonal anti-Calnexin (PA5-19169, Invitrogen), rat monoclonal anti-Pcsk9 (MA524144, Invitrogen), antibodies. Equal loading was evaluated by staining samples with the mouse monoclonal IgG β-actin (sc-81178, Santa Cruz Biotechnology) or mouse monoclonal anti-beta tubulin (MA516308, Invitrogen) antibodies. The proteins were detected with horse radish peroxidase (HRP)-conjugated rabbit anti-goat IgG (H+L) (R21459, Invitrogen), goat anti-mouse IgG, IgM (H+L) (31444, Invitrogen), goat anti-rabbit IgG (H+L) (A27036, Invitrogen) or goat ant-rat IgG (H+L) (31470, Invitrogen) secondary antibodies. The membranes were developed with the SuperSignal™ West Pico PLUS Chemiluminescent Substrate (Thermo Scientific) and imaged by Gel Doc XR+ Gel Documentation System (Bio-Rad).

### Co-culture system

UC-MSCs expressing nucleases were seeded in optimized ratio (L:R nuclease domain or gRNA:Cas9) in a total number of 5×10^4^ cells/well on 12-well plates (Transwell; Corning) and were co-cultured with 5×10^3^ of HEK293T-EGFP or UC-MSC-EGFP cells seeded into 0.4-μm transwell membrane for 8 days. UC-MSC-WT cells served as a negative control for EV-DN production. EGFP-expression was analyzed in the target cells by flow cytometry.

## EVs transfer to recipient cells

EGFP-expressing cells were seeded on 24-well plates (5×10^4^ cells/well) in respective cell culture media and were grown for 24 h to let them adhere. Next, the cells were treated with EVs containing hybrid nucleases (approximately 10^7^ EVs/10^3^ cells) for indicated time points, after which the cells were washed twice with PBS and collected for analysis. To analyze transcripts levels for the nucleases, the recipient cells were collected for RNA extraction at 2, 6, 16 and 26 h of co-incubation. For multiple treatments with EVs, the cells were co-incubated with EVs for 24 h, washed with PBS and after another 6 h new EVs were added, up to five times. 5 days after the last treatment, the cells were collected for FACS analysis for EGFP expression.

### RNA extraction and real time qPCR

Total RNA was extracted from cells using the GeneMATRIX Universal RNA Purification Kit (Eurx), including DNAse I digestion (Ambion) to remove DNA. To isolate RNA from EVs, the Total Exosome RNA and Protein Isolation Kit (Invitrogen) was used. The obtained RNA was reverse transcribed into cDNA using the NG dART RT Kit (Eurx). Transcript levels were measured using the real-time PCR method with the SYBR Green Master Mix (Applied Biosystems) and specific primer pairs listed in the Supplementary Table 7. Quantification of mRNA content was performed on the 7500Fast Real-Time PCR System (Applied biosystems) using the ΔΔCt method with β-2-microglobulin as endogenous control.

### In vivo delivery of EVs

All the animal studies were carried out according to guidelines and protocols approved by the II Local Ethical Committee in Krakow (agreement no #92/2014 and #203/2018). The NOD/SCID-EGFP (NOD.Cg-Prkdc<scid>Il2rg<tm1Wjl>Tg(CAG-EGFP)1Osb/SzJ) mice were purchased from the Jackson Laboratories (JAX# 021937). The experiments were performed using 8-weeks old animals, which were injected intravenously or intraperitoneally with 10^10^ EVs containing nuclease gene editing system (ZFN, TALEN or CRISPR/Cas9) targeting either EGFP or mouse *Pcsk9* gene. Control animals received the same amount of WT-EVs or PBS (in 100 µL volume). Mice were sacrificed 7 days post-injection, selected tissues (heart, lungs, spleen, liver and kidneys) were isolated, mechanically disrupted by passing through a filter with 70 µm pores to obtain single cell suspension and were used for analysis. To detect EGFP gene knockout, the samples were fixed in 2% PFA and analyzed by flow cytometry using the BD LSRFortessa (BD Biosciences). Selected samples with high level of EGFP gene knockout were used for genomic DNA extraction and subsequent genetic analyses.

Liver samples from Pcsk9-2-EVs treated animals were subjected to gDNA, RNA and protein isolation from the same samples using the GeneMATRIX DNA/RNA/Protein Purification [Eurx].

### T7E1 assay

Detection of the CRISPR/Cas9-induced mutations in mouse *Pcsk9* gene was performed with the EnGen Mutation Detection Kit (New England BioLabs; NEB). PCR flanking the CRISPR target site was PCR amplified and products were purified using the GeneMATRIX PCR/DNA Clean-Up Purification Kit (Eurx). 120 ng of the purified PCR product resuspended in 12 μl of buffer 2.1 (NEB) was denatured at 95°C for 5 min and subjected to re-annealing at 95–85°C with ramping at −2°C/s and 85–25°C at −0.1°C/s. Next, the products were treated with T7 Endonuclease I at 37°C for 15 min and inactivated by treatment with Proteinase K at 37°C for 5 min. Untreated samples served as controls. Digested and undigested samples were visualized in 2% agarose gel (Eurx) stained with the SimplySafe nucleic acids stain (Eurx) with the Gel Doc XR+ gel imaging system (Bio-Rad). Quantification was determined by measuring the ratio of cleaved to uncleaved substrate after background subtraction.

### Sequence analysis

Nuclease-edited HEK-EGFP cells were expanded as single clones and lysed with the Direct PCR Lysis Reagent Cell (PeqLab). 2 µl of the lysate was used for PCR-based amplification of the target locus with the Phire Hot Start II DNA Polymerase (Thermo Scientific) using specific primer pairs: forward: 5’-CTACGGCAAGCTGACCCTGAA-3’ and reverse: 5’-GAACTCCAGCAGGACCATGT-3’. The obtained amplicons were purified using the GeneMATRIX PCR/DNA Clean-Up Purification Kit (Eurx) and sequences were analyzed by Sanger sequencing (Genomed) with the GFP-R2-qPCR primer: 5’-TCTCGTTGGGGTCTTTGCTC-3’. The sequences were aligned and visualized using the Clustal Muscle bioinformatics tool [24].

Indels analysis by next generation sequencing was performed on genomic DNA extracted from mouse tissues, using the GeneMATRIX gDNA Isolation Kit (Eurx). Short DNA fragments (270-280 bp) containing the genome-edited target site were PCR-amplified with specific primers (core sequences: forward: 5’-GAAGGGCATCGACTTCAAGG-3’ and reverse: 5’-ATGTGATCGCGCTTCTCGT-3’ for ZFN, or forward: 5’-GGCAAGCTGACCCTGAAGTT-3’ and reverse: 5’-AAGTCGATGCCCTTCAGCTC-3’ for TALENs and the CRISPR/Cas9 targeting the EGFP gene, or forward: 5’-CGTCCATGTCCTTCCCGAG-3’ and reverse: 5’-ACCCATACCTTGGAGCAACG-3’ for mouse *Pcsk9* gene) containing unique 8-mer barcodes and Illumina adapters (synthesized by Genomed). The amplicons were purified using the GeneMATRIX PCR/DNA Clean-Up Purification Kit (Eurx) and were subjected to library preparation with the Nextera XT Kit (Illumina). Libraries were sequenced using v2 Illumina Sequencing Kit on MiSeq platform (Illumina) as pair-end reads (2×250 bp). Data pre-processing was executed using MiSeq Reporter (MRS) v2.6 and contained automated samples demultiplexing and generating Fastq files. Next, adapter sequences were removed (Cutadapt) and sequences with poor quality were discarded from analysis (quality <25, minimal length 150 bp, Cutadapt). Single reads were separated based on internal sequence identifiers using Ea-utils (Genomed). The sequencing data were then further aligned to the respective reference sequences (EGFP and Mus_musculus - GRCm38 for *Pcsk9*) with the use of Subread v2.0.0 read aligner [25] and the settings allowing to discover genomic mutations including short indels and structural variants. The short indels as well as larger rearrangements were stored in the Variant Call Format (VCF). These were then processed with home-made R [26] script in order to identify frequent indels of interest.

### Statistical analysis

All the experiments were performed at least twice in duplicate. The data is presented as means ± standard deviations (SD) or as indicated. Statistical analyses were done with Student’s t-tests, calculated using Microsoft Excel or one-way ANOVA and Tukey’s multiple comparisons test using GraphPad Prism7 (GraphPad Software, La Jolla, CA, USA). *p* values of less than 0.05 were considered statistically significant (* p<0.05; ^$^ p<0.01; ^#^ p<0.001).

## Results

### Generation of UC-MSC lines stably expressing designer nucleases

In our proof-of-concept study, we aimed to test the efficacy of EVs in delivering DNs to recipient cells. We wanted to directly compare the effectiveness of the three nucleases systems: ZFN, TALEN and CRISPR/Cas9, upon EV-mediated delivery. We selected the *EGFP* as a target gene to easily monitor gene knockout efficiency by flow cytometry, since the decline in EGFP expression should correspond to the frequency of modified alleles. We chose UC-MSCs as the producer cells expressing these nucleases, because these are the least immunogenic cells known so far [27]. Thus, EVs collected from these cells can potentially be used in allogeneic settings, without the concern of triggering immune response. We isolated MSCs from umbilical cord using an explant method as previously described [17]. The obtained cells exhibited typical features of MSCs, including spindle-like shape, expression of surface markers: CD166, CD73, CD90, CD44, CD105 and lack of expression of hematopoietic antigens: CD34, CD45, CD14, CD16 and HLA-DR (Supplementary Figure 1A). They also differentiated into osteo-, adipo-, and chondrocytes *in vitro* (Supplementary Figure 1B).

To obtain a long-lasting source of EVs containing DNs and to reduce the risks associated with constitutive nuclease expression, we generated isogenic UC-MSC lines stably expressing single nuclease components (Figure 1). For this purpose, we constructed a series of DN expression vectors containing single domains of each nuclease system, along with puromycin resistance gene, for subsequent selection of genetically modified cells (Figure 1A). We used previously validated nucleases, with proven high EGFP gene knockout activities (Figure 1B) [3,28,29]. We established UC-MSC-DN cell lines which was confirmed on genomic level by PCR-based genotyping using specific primer pairs (Figure 1C, Supplementary Table 1). Next, we verified whether UC-MSC-DNs express the nucleases at mRNA and protein levels. Real-time quantitative PCR (qPCR) revealed that transcript levels for DNs were at significantly higher level in UC-MSC-DNs, in comparison to unmodified cells (Figure 1D). In case of ZFNs, we found that ZFN-R was expressed on a higher level than ZFN-L (Figure 1D, left). Similarly, we found gRNA on a higher level in comparison to *Cas9* (Figure1D, right). The differences resulted most likely from different integration patterns. TALEN-L and -R mRNA levels were similar (Figure 1D, middle). Along with mRNA for DNs, we measured relative expression levels for genes responsible for the maintenance of multipotency (*NANOG, SOX2*), regulators of proliferation (*c-MYC*) and apoptosis (*BAX, BCL-2*) and markers of mesenchymal lineage (*COL1A1, MSX1*). We observed, that ZFN integration upregulated *MSX1* (Figure 1B, left), whereas TALENs integration led to increased expression of multipotency markers *NANOG, SOX2*, the mesenchymal gene *MSX1* and the anti-apoptotic gene *BCL-2* (Figure 1D, middle). In case of gRNA-expressing cells, we found upregulated *NANOG*, whereas Cas9-expressing cells displayed elevated transcript level for *SOX2, MSX1* and *BCL-2* (Figure 1D, right). Importantly, all the nucleases were also expressed at protein level in UC-MSC-DNs, as verified by Western blot analysis (Figure 1E).

**Figure 1.**
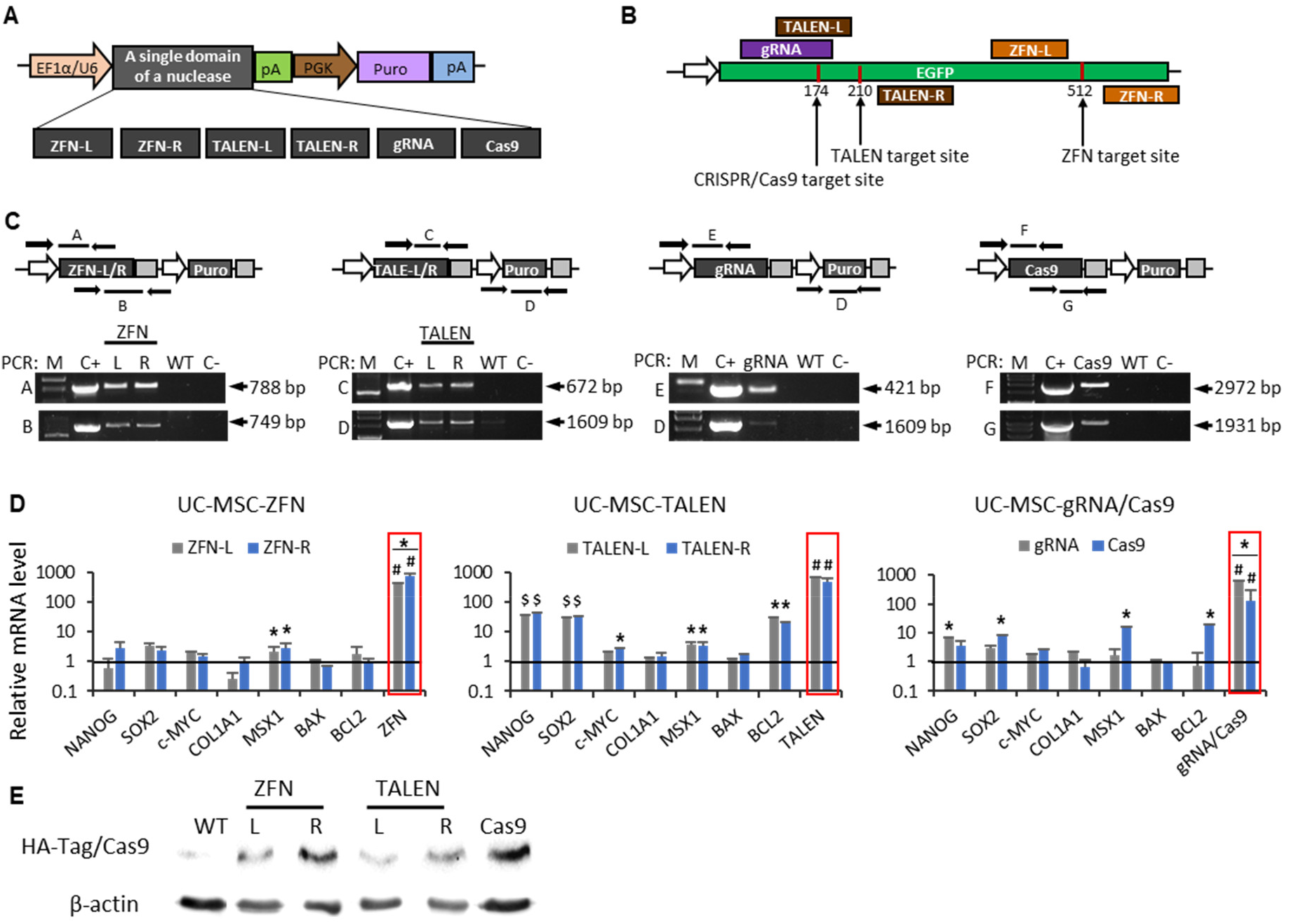
Characterization of UC-MSC lines stably expressing single domains of ZFNs, TALENs and the CRISPR/Cas9 system targeting EGFP gene. **A**. A scheme of plasmids expressing designer nucleases. Block arrows indicate gene promoters and rectangles indicate genes or regulatory elements. Core elements: EF1α - promoter for elongation factor 1 alpha; U6 – U6 nuclear promoter for small RNA expression by polymerase III; pA-polyadenylation signal; PGK – phosphoglycerate kinase 1 promoter; Puro – puromycin resistance gene. **B**. A cartoon showing the binding and the target sites for ZFNs, TALENs and the CRISPR/Cas9 system in the EGFP gene. **C**. Analysis of genomic DNA (gDNA) of UC-MSCs stably expressing designer nucleases (UC-MSC-DN). Upper part: schematic representation of nucleases sequences. Black arrows illustrate primers binding sites for PCR-based genotyping. For each nuclease two reactions were performed, indicated by upper-case letters. Lower part: results for genotyping visualized by agarose-gel electrophoresis. **D**. Relative gene expression levels in UC-MSC-DN. Genes related to apoptosis (*BAX, BCL2*), mesoderm (*COL1A1, MSX1*), ectoderm (*PAX6*), endoderm (*SOX17*), pluripotency (*NANOG, SOX2*) proliferation status *(c-MYC*) and nucleases were studied. Data were normalized to the expression of an endogenous gene (β2-microglobulin) and compered to results obtained from wild type UC-MSC. Value equal to 1 represents gene expression in UC-MSC-WT. The graphs present the mean ± SD from three experiments run in duplicate. Statistical significance at p<0.05, p<0.01 and p<0.001 is indicated with *,^$^ or ^#^, respectively **E**. Protein expression of DN in genetically modified UC-MSC. Antibodies directed toward HA-epitope or Cas9 were used to detect expression of designer nucleases in UC-MSC-DN. Anti-β-actin antibody was used as control.

To further investigate the safety profile of UC-MSC-DNs, we performed real-time qPCR analysis of a series of genes involved in immunomodulatory activity of MSC. Among thirteen molecules tested, we found only macrophage inflammatory protein 1 - alpha (*MIP1-α*) to be significantly upregulated in all UC-MSC-DNs (Supplementary Figure 2). UC-MSC-ZFNs also exhibited higher expression level of interleukin 8 (*IL-8*), UC-MSC-TALENs for indoleamine-pyrrole 2,3-dioxygenase (*IDO*), UC-MSC-gRNA/Cas9 for *IDO* and *IL-8*, and UC-MSC-gRNA additionally for tumor necrosis factor alpha (*TNFα*) (Supplementary Figure 2). Other analyzed factors, including *COX1, COX2, IL1β, MIF, NOTCH2, TGFβ, TLR3, IL-6* remained unchanged (Supplementary Figure 2).

### Isolation of EVs enriched in designer nucleases

Having confirmed that UC-MSC-DNs produce DNs as mRNAs and proteins, we scrutinized the content of EVs isolated from conditioned media (CM) collected from these cells (DN-EVs: ZFN-EVs, TALEN-EVs, CRSPR/Cas9-EVs). To avoid contamination with FBS-derived EVs, we prepared CM by culturing sub-confluent UC-MSC-DNs in serum-free media, containing 0.5% bovine serum album (BSA) for 48 h. CM from standard serum-based cell culture condition served as control. EVs were isolated using sequential ultracentrifugation method at increasing centrifugal speeds, as previously described [17]. The obtained EVs were thoroughly characterized by multiple tests, to meet the criteria launched by the International Society for Extracellular Vesicles (ISEV) in 2018 [15]. First, we measured the size and the concentration of EVs by NanoSight. The analysis revealed that the majority of EVs had approximately 99-110 nm in diameter with no significant differences between the analyzed EV types (Figure 2A, Supplementary Figure 3, Supplementary Table 2). Isolation procedure from 90 mL of CM yielded in approximately 6-12×10^10^ EV particles (Supplementary Table 2), composed of large and small vesicles, as demonstrated by transmission electron microscopy (TEM) (Figure 2B, Supplementary Figure 4). The EVs displayed the presence of exosomal markers (CD81, CD9, Flotillin-1, Syntenin) and did not contain Calnexin, a protein associated with endoplasmic reticulum, as revealed by Western blot analysis (Figure 2C). The vesicles also contained proteins encoding DNs (Figure 2C). Further characterization revealed that EVs shared common surface antigens with their parental cells, including expression of CD90, CD105, CD309, CD147 and lack of expression of hematopoietic marker CD45, which was analyzed by high-resolution flow cytometry (Figure 2D). Noteworthy, the EVs were highly enriched in transcripts encoding DNs in comparison to unmodified (wild type; WT) EVs (Figure 2E). However, we observed differences in mRNA concentration for individual components of DNs. With this respect, ZFN-R-EVs contained higher level of mRNA encoding ZFN, in comparison to ZFN-L-EVs. Similarly, gRNA transcripts were at higher level than Cas9 transcripts in EVs. In turn, mRNA levels for L and R TALEN domains were similar in both EV types (Figure 2E), which correlated to the DNs expression pattern found in producer cells. Irrespective of the nuclease type, concentration of transcripts for DNs was significantly higher in DN-EVs collected in serum-free media in comparison to standard serum-based conditions (Figure 2F), which is of great significance for future clinical applications to treat human diseases.

**Figure 2.**
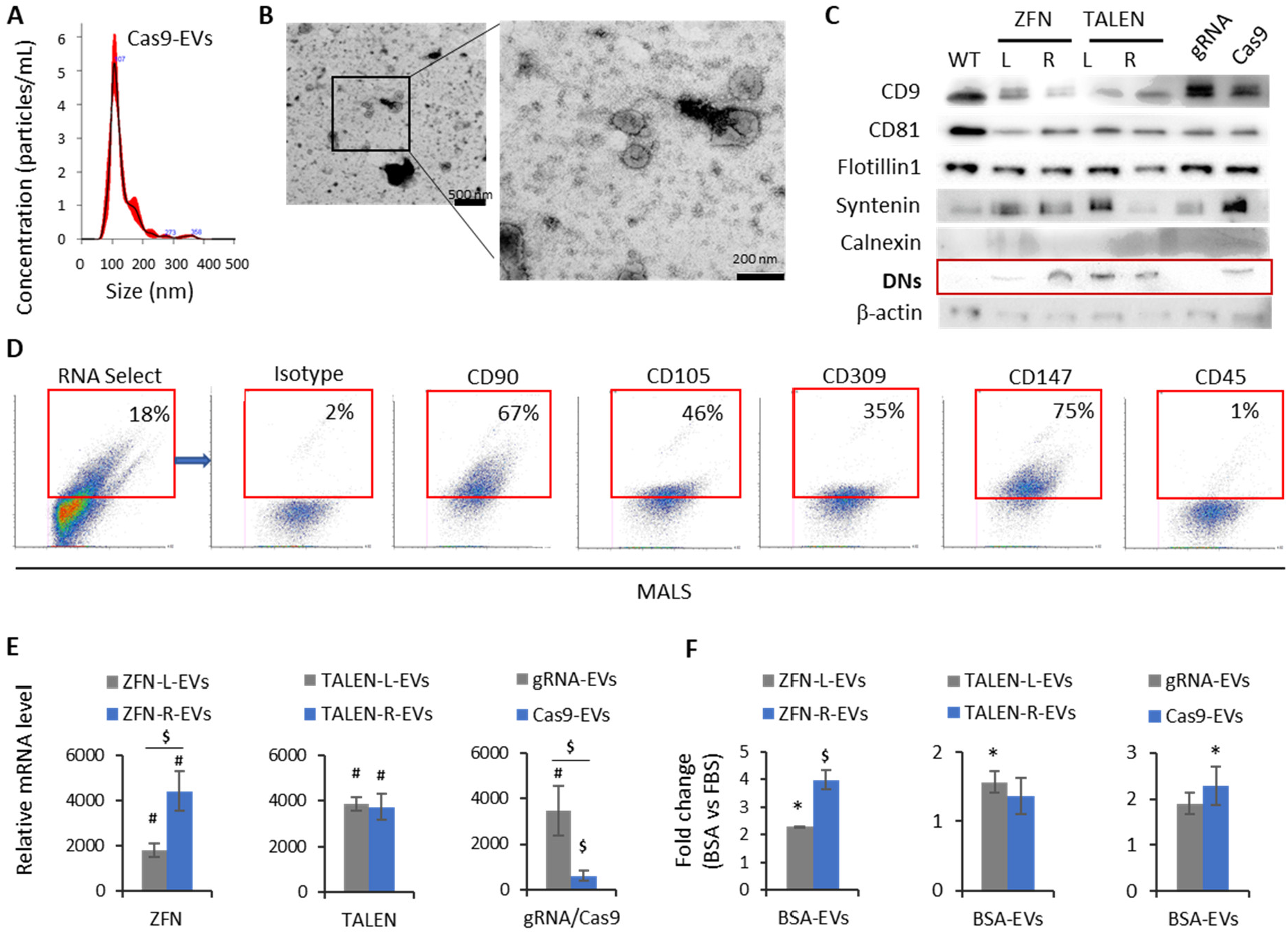
Characterization of EVs derived from UC-MSC-DNs. **A**. Size distribution of DN-EVs analyzed by the NanoSight NS300 instrument. Representative histogram for UC-MSC-Cas9-EVs is shown. **B**. Visualization of DN-EVs using Transmission Electron Microscopy (TEM). Representative TEM pictures for UC-MSC-Cas9-EVs are shown with a scale bar of 500 nm and 200 nm. **C**. Western blot analysis for EV markers, including CD9, CD81, Flotillin-1, Syntenin (positive markers) and ER-associated Calnexin (negative marker). ZFNs and TALENs were detected using anti HA-tag antibody, and Cas9 was detected with anti-Cas9 antibody. Equal loading was confirmed by detection of beta actin. **D**. Surface antigen analysis on UC-MSC-EVs using A50-Micro Flow Cytometer (Apogee Flow Systems). The particles were first gated for SYTO RNASelect-positive events, then the expression of indicated surface proteins was measured. MALS – Medium Angle Light Scatter, corresponding to the particles size. **E**. Analysis of mRNA levels for designer nucleases (ZFN, TALEN and gRNA/Cas9) in DN-EVs using real time qPCR technique and calculated with the ΔΔCt method. Expression of β2-microglobulin was used as endogenous control and mRNA expression levels were calibrated with WT-EVs (level 1 on y axis). Results are presented as the mean ± SD from three experiments performed in duplicate. Statistical significance at p<0.01 and p<0.001 is indicated with ^$^ or ^#^, respectively. **F**. Comparison of DNs mRNA levels in UC-MSC-DN-EVs collected in serum-based medium to the mRNA expression levels obtained in EVs collected in serum-free medium, containing 0.5% BSA. Expression of β2-microglobulin was used as endogenous control and FBS-EVs as calibrator (level 1 on y axis). The graphs show the mean ± SD from two experiments performed in duplicate. Statistical significance at p<0.05 and p<0.01 is indicated with * or ^$^, respectively.

### Knock-out of EGFP gene mediated by DN-EVs *in vitro*

In the next step we used UC-MSC-DN-EVs to achieve EGFP gene knockout in target cells, measured as the decline in fluorescence signal (Figure 3A). For this purpose, we generated EGFP-expressing cell lines, by lentiviral transduction of human embryonic kidney cell line - HEK293T and primary cells - UC-MSCs, using low multiplicity of infection (MOI). We established HEK293T-EGFP cell line (here marked as HEK-EGFP) containing a single copy of EGFP gene, by seeding the cells in limiting dilution and copy number validation in single cell-derived clones by Copy Caller Software (Supplementary Figure 5A). In turn, we obtained polyclonal UC-MSC-EGFP cell population expressing EGFP gene at low level upon sorting out EGFP^dim^ cells. We confirmed stable integration of the transgenic cassette in these cells by PCR-based genotyping and expression of EGFP gene by real-time qPCR and Western blot methods (Supplementary Figure 5B).

**Figure 3.**
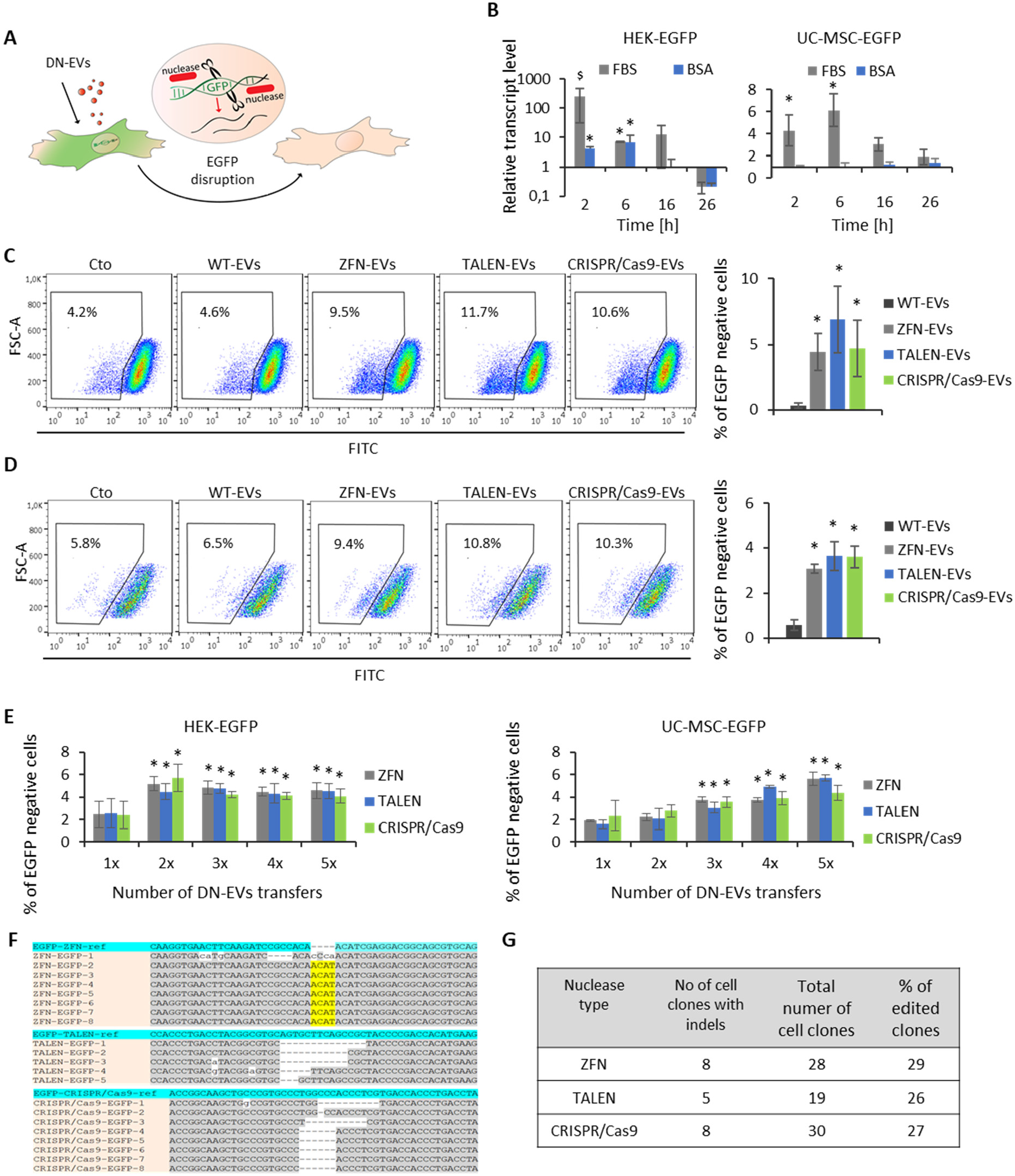
EGFP knockout *in vitro* in EGFP-expressing cell lines mediated by DN-EVs. **A**. A cartoon showing the principle of the experiments. DN-EVs activity in target cells should lead to the disruption of EGFP gene and the decline in EGFP protein expression. **B**. Transcript levels for ZFNs in HEK-293T-EGFP and UC-MSC-EGFP cells after incubation with ZFN-EVs at indicated time points (2, 6, 16 and 26 h). The target cells were cultured in standard condition in serum-containing medium (FBS) or were synchronized in cell cycle by starvation in serum-free medium, containing 0.5% bovine serum albumin (BSA). **C, D**. Co-culture system of DN-expressing cell lines with HEK-EGFP (**C**) or UC-MSC-EGFP (**D**) cell lines. DN-EVs-producer cells and target cells were cultured in Boyden chambers separated by a mesh filter with 0.4 µm pores for 8 days. EGFP knockout was measured by flow cytometry. Representative dot plots are shown (FlowJo). The graphs present the mean + SD from three experiments. Statistical significance at p<0.05 is indicated with *. **E**. Transfer of DN-EVs to HEK-EGFP and UC-MSC-EGFP cell lines. Cells were treated up to five times with DN-EVs in a dose of 10^7^ EVs/10^3^ cells. EGFP knockout was measured at day 5 after the last EVs transfer by flow cytometry. Results are presented as the mean + SD from three experiments performed in duplicate. Statistical significance at p<0.05 is indicated with *. **F**. Analysis of EGFP gene disruption in single cell-derived clones of HEK-EGFP cells upon transfer of DN-EVs. Genomic DNA was extracted and used as a template to amplify regions flanking the DN site by PCR. The obtained amplicons were purified and subjected to sequence analysis by Sanger method. The identified indels were aligned to the WT sequence and visualized with the Clustal Muscle bioinformatics tool. **G**. Calculation of indels frequency in HEK-EGFP single cell-derived clones treated with DN-EVs.

We evaluated DN-EVs-mediated knockout efficiency in target cells in two ways: i) in a co-culture system and ii) by direct DN-EVs transfer. However, knowing that ZFN-L and R domains, as well as gRNA and Cas9 were not expressed at equal levels in DN-EVs, first, we optimized the ratio of single components of these nucleases for efficient EGFP gene knockout. For this purpose, we co-incubated HEK-EGFP cells with DN-EV pair containing ZFN and the CRISPR/Cas9 system at various ratios and with repeated doses. We achieved almost 6% of EGFP gene knockout for ZFN at 2:1 ratio of ZFN-L to -R EVs and over 6% EGFP knockout for the CRISPR/Cas9 system at 5:1 ratio of gRNA to Cas9-EVs (Supplementary Figure S6). These ratios were selected for further experiments. EVs containing TALEN-L and R domains were used in 1:1 ratio.

To further optimize our genome editing system, we tested whether cell cycle synchronization improves DNs transfer from EVs to target cells. This analysis was limited to the ZFN-EVs. The target cells were either cultured in standard conditions (cell culture medium supplemented with 10% FBS) or starved for 24h (cell culture medium containing 0.5% BSA) before the addition of ZFN-EVs. Next, we measured ZFN transcript levels in HEK-EGFP and UC-MSC-EGFP cells at indicated time points using real-time qPCR method. The results demonstrate that mRNA for ZFNs was detectable in the recipient cells, particularly after 2h upon DN-EVs addition (Figure 3B). The EVs uptake was more efficient in cells cultured in standard cell culture medium, which were not starved (Figure 3B) and such conditions were used in further experiments.

To confirm that direct DN-EVs transfer from producer to recipient cells is responsible for genome modification, we used a co-culture system in a Boyden chamber. The separating membrane contained pores with 0.4 µm in diameter, which prevented cell-to-cell interactions and ensured that only EVs could translocate. We seeded UC-MSC-DN cells in the bottom of the wells, to ensure abundant production of DN-EVs, and the EGFP-expressing cells on the upper part of the transwell. The cells were co-cultured for 8 days, which was the end-point for the measurement of EGFP expression. According to the results and after background subtraction, we obtained up to 7% of EGFP gene knockout in the co-culture system with HEK-EGFP cells (Fig. 3C) or almost 4% in UC-MSC-EGFP cells (Figure 3D). There were no significant differences between various nucleases types.

Next, we tested direct DN-EVs transfer to recipient cells. For this purpose, we added either ZFN-EVs or TALEN-EVs or CRISPR/Cas9-EVs to the HEK-EGP or UC-MSC-EGFP cells, respectively, and we measured reduction in EGFP gene expression by flow cytometry. To potentially enhance the results, we treated the cells with DN-EVs up to five times, with 24 h interval. Ultimately, we obtained almost 6% of EGFP gene knockout in both HEK-EGFP and UC-MSC-EGFP cells, irrespective of the nuclease type (Figure 3E). Interestingly, the highest knockout rate in HEK-EGFP cell was achieved after second DN-EV transfer and further treatment did not improve the outcome (Figure 3E, left). In contrast, repeated doses of DN-EVs increased gene knockout efficiency in UC-MSC-EGFP cells (Figure 3E, right).

Next, we wanted to confirm that decline in EGFP expression in HEK-EGFP gene truly resulted from DN-EV-mediated gene disruption. Therefore, we analyzed genomic DNA (gDNA) of EGFP negative single cell-derived clones after direct transfer of DN-EVs by Sanger sequencing. We identified 8, 5 and 8 cell clones with indels upon ZFN, TALEN or CRISPR/Cas9-EVs treatment, out of 28, 19 and 30 analyzed cell clones, respectively (Figure 3F,G). In case of ZFN-EVs, we observed predominantly insertion of 4 nucleotides (Figure 3F, upper panel). Whereas activity of TALEN-EVs and CRISPR/Cas9-EVs led to deletions of 1 to 14 nucleotides (Figure 3F, middle and lower panels, respectively). Taken together, we detected gene knockout in 26 to 29% of the analyzed cell clones (Figure 3G). A relatively high percentage of EGFP negative cells with a silenced transgene generated high background of unedited cells. Genome editing efficiencies were similar for all three nucleases systems.

### Efficient *in vivo* delivery of DNs by EVs

After *in vitro* studies, we validated our DN-EVs delivery system *in vivo*. For this reason, we transplanted DN-EVs targeting the *EGFP* gene to NOD/SCID-EGFP mice (Figure 4A). We tested two routes of EVs administration: i) intravenous (*iv*; into the *venous sinus*) and ii) intraperitoneal. We also tested different doses of DN-EVs, to find optimal gene targeting conditions. After 7 days post-injection, the mice were sacrificed and we collected selected organs, including spleen, lungs, heart, kidneys and liver for analysis. To release single cells, we minced the tissues mechanically and passed the cells through cell filters with 70 µm pores. Subsequently, we measured EGFP gene expression by flow cytometry and calculated the knockout rates after background subtraction from PBS-injected animals. The results show that single intravenous administration of 10^10^ DN-EVs resulted in 1 to 15% of EGFP-negative cells, with the highest percentage of gene disruption in the liver (Figure 4B). We have not observed significant differences between different EV types, containing either ZFN, TALEN or the CRISPR/Cas9 system. Multiple injections of DN-EVs (here: CRISPR/Cas9-EVs) increased the fraction of EGFP-negative cells up to 30% in the liver (Figure 4C). Interestingly, intraperitoneal transplantation of a single dose of 10^10^ DN-EVs resulted in higher EGFP-gene knockout rate in several organs, reaching up to 50% gene knockout in the liver (Figure 4D). This way of DN-EVs administration was more effective for biological activity of DN-EVs in the kidneys, in comparison to *iv* injection.

**Figure 4.**
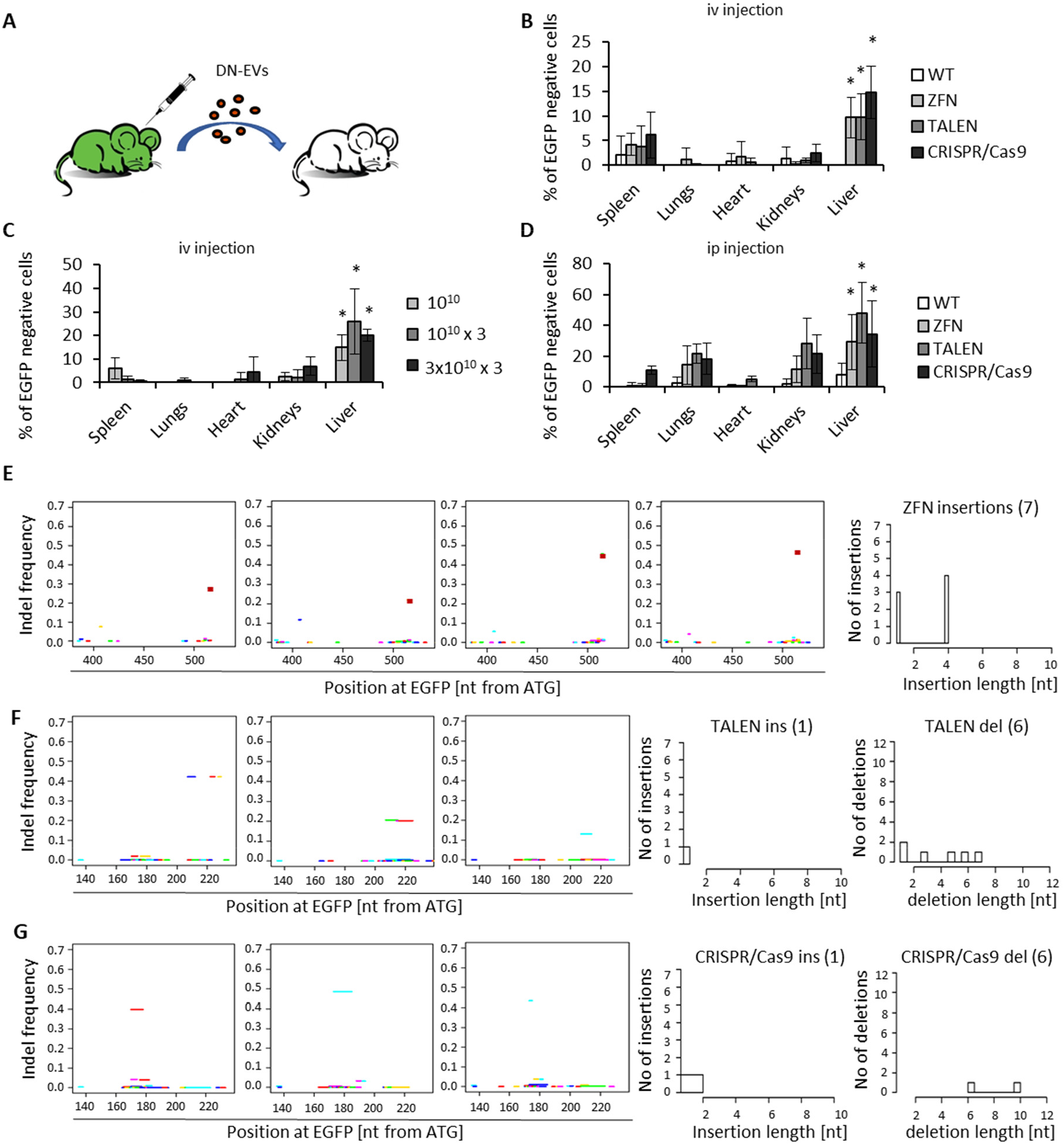
Analysis of EGFP gene knockout in vivo mediated by UC-MSC-DN-EVs. **A**. A scheme showing the principle of the *in vivo* genome editing experiments. Injected stem cells-derived EVs carrying hybrid nucleases targeting EGFP gene, will lead to inactivation of EGFP gene expression and loss of fluorescence. **B**,**C**,**D**. NOD/SCID-EGFP mice (n=5-6 per group) were transplanted intravenously with a single dose of 10^10^ DN-EVs (**B**) or with repeated doses and increasing amount of CRISPR/Cas9-EVs (**C**), or with a single dose of DN-EVs intraperitoneally (**D**). After 7 days post-injection the mice were sacrificed and cells were isolated from various organs (spleen, lungs, heart, kidney, liver) by mechanical disruption. EGFP knockout was measured by flow cytometry. Results are shown as the mean + SD after background subtraction from PBS-injected animals. Statistical significance at p<0.05 is indicated with *. **E**,**F**,**G**. Sequence analysis of liver samples injected with DN-EVs (n=3-4 per group) using deep sequencing on Illumina platform. A short (∼270 nt) region flanking the nucleases target sites were PCR-amplified, purified and sequenced. The plots are showing indels frequency in individual animals at the indicated positions from the EGFP gene START site and indels characteristics for a group with respect to indel length and the number. **E**. Indels frequency and characteristics in 4 liver samples from mice treated with ZFN-EVs. **F**. Indels frequency and characteristics in 3 liver samples from mice treated with TALEN-EVs. **G**. Indels frequency and characteristics in 3 liver samples from mice treated with CRISPR/Cas9-EVs.

To confirm EGFP gene disruption at genomic level, we PCR-amplified short fragments of gDNA flanking the target region of interest, in 3-4 liver samples obtained from animals treated with DN-EVs. The amplicons were then subjected to deep sequencing using Illumina platform. The analysis confirmed high editing efficiencies (Figure 4E-G, Supplementary Table 3), corresponding to the knockout frequencies of the flow cytometric measurements. In case of ZFN-EVs, the most dominant sequence change in EGFP gene was identified in a frequency of 0.23 to 0.47 of the total reads and resulted from 1 or 4 nucleotide (nt) insertions (Figure 4E). In turn, TALEN-EVs-treated samples contained modified alleles in a frequency of 0.13-0.42, which was supported by identification of a 1 nt insertion and several deletions of 1-7 nt (Figure 4F). CRISPR/Cas9-EVs activity led to permanent disruption of EGFP gene in a frequency up to 0.48, marked by a 2 nt insertion or deletions of 6 or 10 nt (Figure 4G).

### *Pcsk9* gene knock-out *in vivo* by iPS-DN-EVs

Encouraged by the results from *in vivo* experiments, we wanted to further explore the utility of our EV-based genome editing system towards EVs derived from other stem cell sources. Being at the beginning of the era of personalized medicine, in which utilization of patient-derived induced pluripotent stem cells (iPSCs) is becoming more and more desirable, we tested iPSC-derived EVs as DN delivery vehicles. In this part of research, we focused on the CRISPR/Cas9 platform, since it is highly versatile and easy to customize. To simplify the system, we first checked whether overexpression of a complete genome editing system in one parental cell line is as safe and efficient in the context of DN-EVs production, as overexpressing single components separately in different lines. Using a commercially available hiPSC line (Figure 5A), we generated isogenic cell lines overexpressing either single or both components of the EGFP-targeted system used above. We confirmed stable integration of the expression cassettes in these cells by PCR-based genotyping (Supplementary Figure 7A) and the expression of genome editing components by real time qPCR and Western blot methods, respectively (Supplementary Figure S7B,C). We also compared mRNA levels of genes related to pluripotency (*OCT4, NANOG*), apoptosis (*BAX, BCL2*) and proliferation (*c-MYC, p21*) between genetically modified and unmodified, WT cells (Supplementary Figure S7D). Although we observed downregulation of pluripotency genes in all genetically modified cell populations, the expression of *OCT4* in the gRNA/Cas9-expressing cells remained as high as in parental cells. In turn, *BAX* and *BCL-2* were upregulated in all tested populations, the least in hiPS-Cas9 cell line. In case of cell cycle regulators, we detected the smallest decrease in *c-MYC* expression in hiPS-gRNA/Cas9 cell line, along with unchanged expression of p21. Based on these results, we decided to further proceed with EV-based genome editing system containing both components of the CRISPR/Cas9 system in single EVs.

We aimed to target an endogenous gene, instead of a marker gene, to prove usefulness of our gene editing strategy in a potential therapy of a genetic disease. We chose *Proprotein Convertase Subtilisin/Kexin Type 9* (*Pcsk9*) as a target, since overexpression of this gene has been implicated in hypercholesterolemia [21]. Therefore, reduction of PCSK9 expression mediated by DN-EVs could bring a desirable curative effect. We selected two guide RNAs targeting exon 1 of mouse *Pcsk9* gene at two distinct locations (Figure 5B). After *in vitro* synthesis of the guides, we complexed them with commercially available Cas9 protein and evaluated their activity *in vitro*, using T7 Endonuclease 1 assay. Both guides showed high activity, however, gRNA2 was more active than gRNA1, giving rise to 54% of alleles with indels (Figure 5C). Nonetheless, we used both gRNAs to prepare novel hiPSC lines producing DN-EVs. gRNAs were first cloned into a lentiviral (LV) vector containing Cas9-expression cassette and puromycin resistance gene, and LV particles were used to transduce hiPSCs, to generate stable cell lines. Analysis of gDNA from modified cells confirmed integration of the transgenic cassettes in both, hiPS-Pcsk9-1 and hiPS-Pcsk9-2 cell lines, containing gRNA1 and gRNA2, respectively (Figure 5D). Interestingly, transcript levels for gRNA and Cas9 were slightly higher in hiPS-Pcsk9-1 cell line, in comparison to hiPS-Pcsk9-2 cell line, albeit not significantly (Figure 5E, left). The observed difference negatively correlated with expression of pluripotency markers *OCT4* and *NANOG* in the analyzed cell lines (Figure 5E, right). Importantly, Cas9 protein was readily detectable in both cell lines (Figure 5F).

**Figure 5.**
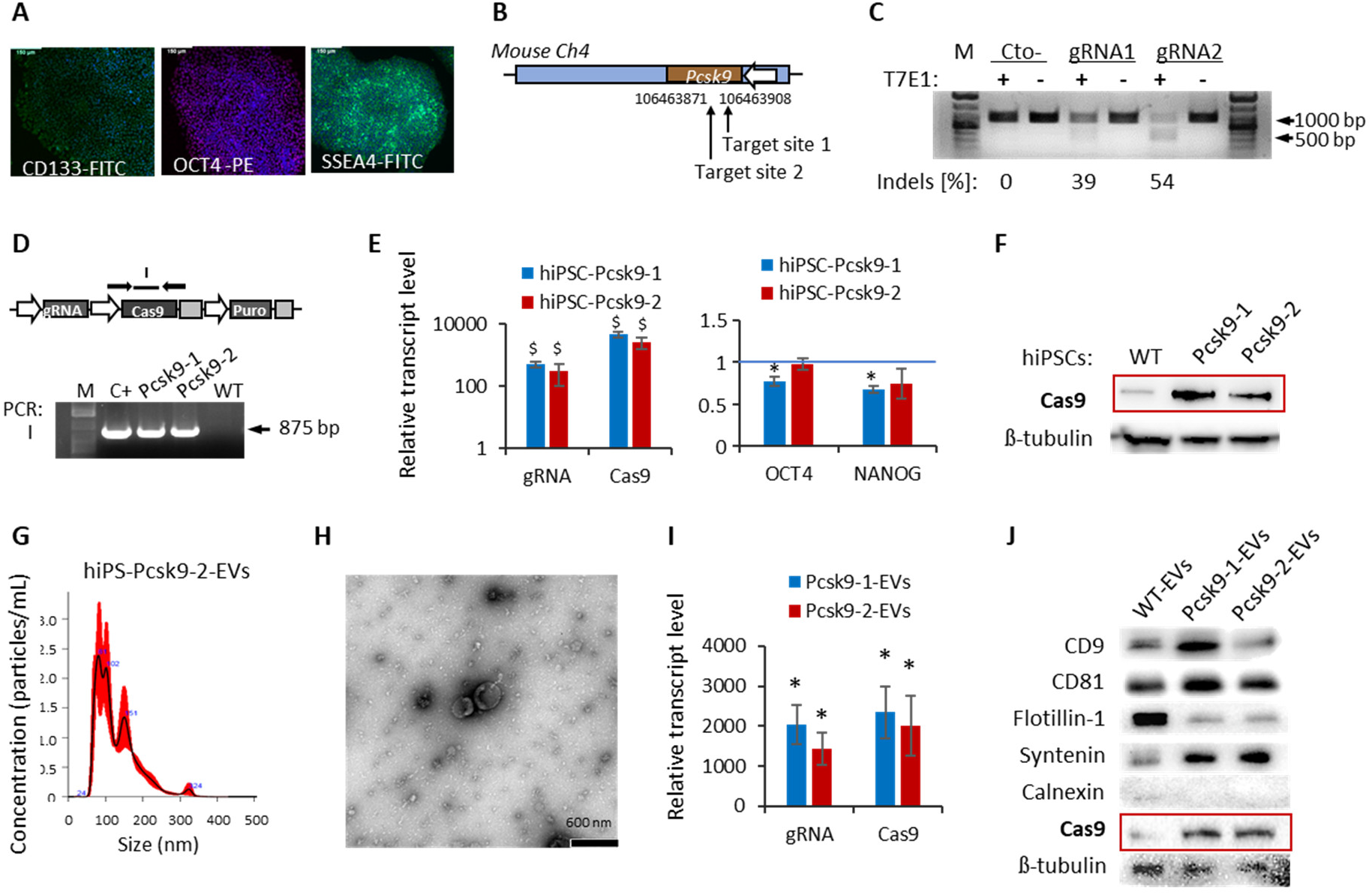
Generation and characterization of hiPSCs and hiPS-EVs expressing the CRISPR/Cas9 system targeting mouse *Psck9* gene. **A**. Detection of pluripotency factors in hiPSC line used in experiments. Representative images of hiPSC colonies stained with fluorescently labeled antibodies against CD133, OCT4, and SSEA-4 from fluorescent microscope. Scale bar is 150 µm. **B**. Localization of target sites on mouse chromosome 4 for two gRNAs targeting *Pcsk9* gene. **C**. Indels frequency for the two selected gRNAs in mouse *Pcsk9* gene measured with the T7 Endonuclease I assay. **D**. Analysis of stable integration of the transgenic cassettes expressing the CRISPR/Cas9 system components in hiPSCs on the gDNA level. Upper part: schematic representation of the expression cassettes. Black arrows illustrate primers binding sites for PCR-based genotyping. Lower part: results for genotyping visualized by agarose-gel electrophoresis. **E**. Transcript levels for gRNA and Cas9 (left) and pluripotency-associated genes: *OCT4* and *NANOG* (right) in genetically modified hiPSCs, relative to control, unmodified cells (level 1 on y axis). Expression of β2-microglobulin was used as endogenous control. The obtained data were calculated with the ΔΔCt method and are presented as the as the mean + SD from two experiments performed in duplicate. Statistical significance at p<0.05 is indicated with *. **F**. Western blot analysis of Cas9 protein expression in hiPS-Pcsk9-1/2 cell lines. Detection of β-tubulin was used as control. **G**. Size analysis of hiPS-Pcsk9-EVs measured with the NanoSight NS300 instrument. Representative histogram for hiPS-Pcsk9-2-EVs is shown. **H**. Representative TEM picture of hiPS-Pcsk9-2-EVs. Scale bar is 600 nm. **I**. Transcript levels for gRNA and Cas9 in hiPS-Pcsk9-EVs, relative to control, unmodified EVs (level 1 on y axis). The data were calculated with the ΔΔCt method using expression of β2-microglobulin as endogenous control. Results are presented as the as the mean + SD from two experiments performed in duplicate. Statistical significance at p<0.05 is indicated with *. **J**. Western blot detection of EVs markers: CD9, CD81, Flotillin-1, Syntenin, Calnexin (negative marker) and Cas9 in hiPSC-Pcsk9-EVs. Equal loading was confirmed by detection of β-tubulin.

As previously, we isolated EV from conditioned media collected from sub-confluent cultures of hiPS-Pcsk9-1/2 cell lines and thoughtfully characterized them. The obtained EV populations were heterogenous, with the mean size of the particles of approximately 130 nm, as measured by NanoSight (Figure 5G, Supplementary Figure 8A, Supplementary Table 4). The EVs were also visualized by TEM (Figure 5H, Supplementary Figure 8B). Next, we measured mRNA levels for gRNA and Cas9 in the EVs by real time qPCR method. We detected both transcripts at significantly upregulated levels, in comparison to EVs from unmodified cells (Figure 5I). Although hiPS-Pcsk9-1-EVs contained gRNA and Cas9 transcript more abundantly than hiPS-Pcsk9-2-EVs, the difference was not statistically significant. Interestingly, gRNA and Cas9 overexpression led to increase in transcript levels for pluripotency genes *OCT4* and *NANOG*, in the EVs (Supplementary Figure 8C). Cas9 protein was also detected in both EV types, as verified by Western blot method (Figure 5J). All the EV types displayed typical EV markers, such as CD9, CD81, Flotillin-1, Syntenin and did not contain Calnexin, as expected (Figure 5J). Considering higher activity of gRNA-2 and not significantly different mRNA levels for the components of the CRISPR/Cas9 system, we selected hiPS-Pcsk9-2-EVs for further studies *in vivo*.

We injected 10^10^ of hiPS-Pcsk9-2-EVs into the venous sinus of 9 experimental animals. After 7 days the mice were sacrificed, the livers were extracted and cells were isolated by mechanical disruption. gDNA was used to identify mutations induced by the CRISPR/Cas9 system at precise genetic locations. For this purpose, we amplified fragments of gDNA flanking the CRISPR/Cas9 target site and analysed the sequences by next generation sequencing. The analysis revealed very high targeting rates, reaching almost 0.5 indel frequency in all experimental animals (Figure 6A, Supplementary Table 5). The most common mutations were deletions of several nucleotides (Figure 6B). A sample picture demonstrating the alignment of the sequenced DNA fragment encompassing deletions generated by the hiPSC-Pcsk9-2-EVs is shown in the Supplementary Figure 9. Notably, the disruption in the mouse *Pcsk9* gene was reflected in decreased expression of its transcript in all examined animals (Figure 6C), as well as in reduced protein level (Figure 6D,E).

**Figure 6.**
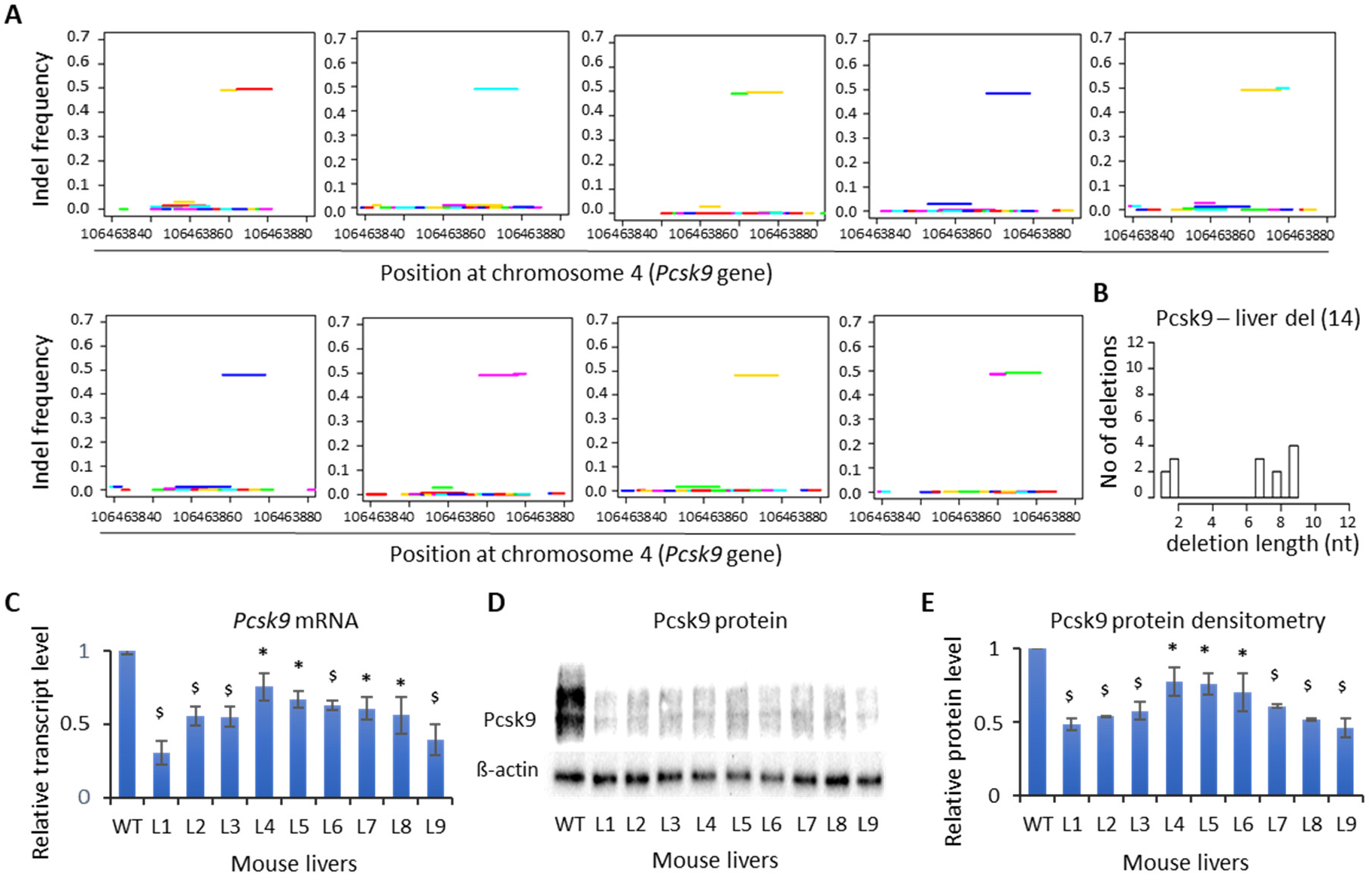
Knockout of *Pcsk9* gene in mouse liver by hiPS-EVs transferring the CRISPR/Cas9 system. Mice (n=9) were injected with hiPS-Pcsk9-2-EVs intravenously and were sacrificed 7 days post-injection. Mouse livers were extracted and cells were isolated by mechanical disruption. Upon gDNA isolation, PCR fragments flanking the CRISPR/Cas9 target site were subjected to deep sequencing using Illumina platform. **A**. Graphs showing indels frequency at indicated genomic position on chromosome 4 of individual animals. **B**. Indels length and number detected in liver samples from all animals. **C**. Transcript level of Pcsk9 gene in liver samples obtained from mouse treated with hiPS-Pcsk9-2-EVs measured with real time qPCR method. The analysis was done using the ΔΔCt method and β2-microglobulin as endogenous control. Level 1 on y axis corresponds to expression level in a mouse liver injected with hiPS-WT-EVs. The data are presented as the as the mean + SD from two experiments performed in duplicate. Statistical significance at p<0.05 or p<0.01 is indicated with * or ^$^, respectively. **D**. Detection of Pcsk9 protein in mouse liver samples obtained from animals injected with hiPS-Pcsk9-2-EVs. Expression of β-actin was used as loading control. **E**. Densitometric analysis of protein level of mouse Pcsk9, relative to β-actin. Results are shown as the mean ± SD from three experiments. Statistical significance at p<0.05 or p<0.01 is indicated with * or ^$^, respectively.

## Discussion

Recent years have brought significant discoveries and meaningful advancements in the field of genome editing, pushing the concept of safe and efficient gene therapy closer to clinical practice. One of the important steps on this path was adaptation of the bacterial defense mechanism, the CRISPR/Cas9 system, to genome editing purposes [4,5]. This system, due to the ease of design, is highly versatile and the most popular nowadays, compared to the previously used technologies. However, in addition to the CRISPR/Cas9, both ZFNs and TALENs have proved their usefulness as medicinal products [30], with the highest number of completed clinical trials using ZFNs (https://clinicaltrials.gov/ct2/home). The key aspects of a successful gene therapy, apart from precision of the genome editing platform, is safety and efficiency of the method applied to deliver therapeutic nucleases into a living organism. Here, we showed highly efficient genome editing *in vivo*, based on the use of EVs derived from human stem cells, carrying ZFN, TALEN and the CRISPR/Cas9 system.

EVs, as natural components of cell-to-cell communication, participate in horizontal transfer of bioactive molecules, including nucleic acids, therefore constitute an attractive technological solution to deliver DNs *in vivo*. Due to their cellular origin, EVs are considered biocompatible, non-toxic, and possess the ability to cross the blood-brain barrier. Thus, they can overcome limitations associated with the current *in vivo* delivery methods of therapeutic nucleases. The efficacy of modified or naive EVs derived from different human cell populations, such as embryonic kidney (HEK293T) cell line [31-34], dendritic cells [35], red blood cells [36] and tumor cells [37,38] has already been demonstrated to deliver small RNAs [31] and the CRISPR/Cas9 system *in vivo* [32,33,36,37]. However, precautious has to be taken when selecting appropriate EV producer cell line for the delivery of a therapeutic cargo. Although tumor cell-derived EVs proved effective in delivering CRISPR/Cas9 into tumors *in vivo* [37], they were also shown to transfer oncogenic elements and endogenous retroviral DNA fragments to healthy cells [39] and to participate in the formation of pre-metastatic niches [40]. Other EV-producer cells also possess their limitations. Blood cells-derived EVs must present a particular repertoire of human leukocyte antigens on their surface, in order to escape immune response upon injection to a living organism. In turn, a popular HEK293 cell line, although approved for production of therapeutic proteins by the European and American regulatory agencies [41], possesses unstable karyotype, tumorigenic potential [42], is amenable to genome alterations in response to genetic manipulations [43]. All these factors may contribute to production of EV containing undesired cargo. With this respect, HEK293 cells-derived exosomes were shown to impact on cadherin-associated signaling pathway *via* their microRNAs (miRNAs) and to increase motility of breast cancer cells [44]. Together, these data indicate that EVs are not inert and along with an intended message, they co-deliver other bio-molecules, which may trigger an unexpected biological response in recipient cells. On the other hand, optimal selection of parental cells may provide EVs with molecular cargo positively impacting on target cells. For this reason, we carefully selected producer cells for harvesting therapeutic EVs. First, we focused on UC-MSCs, then we extended the system towards iPSCs, since both cell types are currently utilized in medical applications.

The most advantageous feature of UC-MSCs is their lack of immunogenicity, hence they can be used in allogeneic therapies in variety of diseases [27]. Similarly, UC-MSC-derived EVs, with their immune-privileged profile and pro-regenerative properties, are gaining more and more popularity as future medicinal products [45]. In turn, hiPSCs opened great perspectives in the field of personalized medicine [46]. However, their tumorigenic potential restricts their clinical applications. Therefore, hiPS-EVs, which contain pro-regenerative bioactive cargo, constitute a competitive solution as a new therapeutic tool [17,19,47]. As we previously demonstrated, intramyocardial injection of mouse iPS-EVs did not trigger formation of teratomas in experimental animals, in the contrary to mouse iPSCs [19]. In the present study, we confirmed that hiPSC-EVs are not capable of generating tumors: up to 90 days after subcutaneous injection we did not detect any tumors (n=5, data not shown).

Importantly, we set up our system in serum-free conditions, to avoid any contamination of serum-derived bio-molecules. We have previously shown that media composition impacts on biological properties and immunomodulatory potential of UC-MSCs and their EVs [18]. Based on these data and in order to advance our genome editing system on potential clinical use, we optimized EVs production in serum-free conditions. Apparently, culturing producer cells in serum-free media yielded in EVs enriched in transcripts for DNs (Figure 2F).

Our *in vivo* biodistribution studies have shown that EVs accumulate mainly in the liver (Figure 4B-D), which is in accordance with the results obtained by others [48]. We took advantage of this natural EV tropism and we showed highly efficient targeting of *Pcsk9* gene in the liver of the experimental animals using our DN-EV platform (Figure 6). These results can be further exploited for therapeutic use of stem cells-derived EVs carrying hybrid nucleases in the treatment of hypercholesterolemia [49]. Importantly, genetic inactivation of *Pcsk9* gene in the animal livers using engineered meganucleases [50], ZFN [51], base editors [52], or chemically improved gRNA/Cas9 complexes [53] resulted in persistent reduction of serum cholesterol, highlighting long-term benefits of a gene therapy approach. Notably, such powerful *in vivo* activity of DN-EVs can be extended towards applications in the treatment of other genetic diseases, which sheds new light on opportunities in medicine of the 21^st^ century.

Regarding safety and efficiency of our *in vivo* genome editing platform utilizing first generation CRISPR/Cas9 system or other hybrid nucleases, we deem that it is possible to enhance both, by introducing certain improvements. For the next generation DN-EVs, we are considering to: i) Restrict the tropism of the EVs towards a certain type of cells or tissues by inserting specific ligands on the surface of DN-EVs. Upon binding to particular receptors, which are present only on a desired cell population, we can increase the tropism and targeting rate towards selected cells. ii) Improve packaging of DNs into EVs. In this proof-of-concept project, we relied on DN export from parental cells stably overexpressing the components of genome editing system to EVs. However, stable transgene integration may disturb cellular homeostasis and may lead to changes in gene expression, as we observed in case of certain transcripts in UC-MSC-DNs (Figure 1D, Supplementary Figure S2) or hiPS-DNs (Supplementary Figure S7D, Figure 5E). For this reason, we aim to implement a controlled packaging of synthetic Cas9 and gRNA into the EVs by electroporation. Recent studies have proven efficacy of this way in uploading DNA [54] and small RNA [36,55] cargo into EVs. Such techniques will enable obtaining an optimal and dose-regulated nuclease amount in EVs and, importantly, will allow the addition of a repair matrix for gene correction. It has to be emphasized that the majority of genetic diseases result from insufficiency in gene expression, due to inactivating mutations. Thus, correction of a disease phenotype will rely on precise repair of the underlying mutation from a repair template, which has to be co-delivered exogenously. iii) Increase the activity and specificity of DNs by using chemically modified nucleases, including gRNAs or other Cas variants incorporated into EVs, which can substantially reduce the nuclease-induced off-targets *in vivo* [56].

In summary, in this work we showed efficient *in vivo* genome editing using distinct stem cells-derived EVs as delivery vehicles for DNs. Compared to other recently described technologies, our system displays several advantages. It allows for transient DNs activity delivered in vivo as RNA and protein, thus reduces harmful side effects of a nuclease. With respect to the CRISPR/Cas9 system, the Cas9-induced off-target mutagenesis constitutes the major concern of clinical translation of this technology [57]. Prolonged Cas9 activity results in a higher risk of DNA damage. On the contrary, RNP delivery of Cas9 and gRNA substantially reduces genotoxicity [58]. In our EV-based system, we rely on protein and RNA delivery, which minimizes off-targets frequency. Further safety precautions encompass various modifications of gRNA sequence [59,60a], which can be easily adapted to our nuclease delivery platform. Importantly, we used EVs derived from clinically relevant stem cell populations, which can be used in allogeneic therapies (UC-MSC-EVs) or in personalized medicine approaches (iPSC-EVs). EVs are stable in long-term storage, thus the system creates the opportunity to prepare an off-the-shelf product for many patients (UC-MSC-EVs) or can be available upon request if repeated doses are needed (UC-MSC-EVs and iPSC-EVs). Our system can be further adapted for the delivery of other targeted modification systems, including transcriptional activators [61], epigenome editors [62], base editors [52] or components of the prime editing system [63]. Taken together, the EV-based system for delivery of DNs presented in this work opens new horizons in the treatment of genetic diseases.

## Acknowledgements

We thank Francesco Gubinelli (Jagiellonian University) for technical support in mouse studies and Prof. Christopher Baum (MHH Hanover) for merit support.

## Author contributions

SB-W conceived the project, obtained the funding, trained the students, set up and performed the *in vitro* experiments with the help of KK the *in vivo* experiments with KK, KN, NB, KK-W, ASz, MT-C, molecular analyses with KK, KN, NB, KK-W, MP and MT-C. EK sorted out the UC-MSC-EGFP^dim^ cells, analyzed EVs on NanoSight and ApoGee. OW performed TEM analysis. PPŁ analyzed the NGS data. DB provided umbilical cords. TC and CM provided plasmids for ZFN, TALEN and CRISPR/Cas9 targeting EGFP gene. CM, ASc, TC, ZM and EZ-S provided merit support. ZM and EZ-S provided financial and infrastructure support. SB-W wrote manuscript. All the authors revised and accepted the manuscript.

## Disclosure of interest

Jagiellonian University and Medical Center - University of Freiburg have filed a patent application for the use of stem cell-derived EVs as carriers of gene modifying enzymes on behalf of the inventors SB-W, EZ-S. and TC. TC has a funded research collaboration with Cellectis. The remaining authors declare no competing interests.

## Funding

This work was supported by the Homing Plus/2013-7/3 project carried out within the Homing Plus programme of the Foundation for Polish Science (FNP) co-financed by the European Union under the European Regional Development Fund, the SONATA12 project: UMO-2016/23/D/NZ3/01310 from the National Science Centre of Poland granted to S.B.-W., the project TEAM-2012/9-6 (FNP), POIG.01.01.02-00-109/09, STRATEGMED 3 (STRATEGMED3/303570/7/NCBR/2017) funded by National Center for Research and Development (NCBR) to E.Z.-S., and the support from the KNOW grant of the FBBB from the Ministry of Science and Higher Education of Poland.

