## Supplementary files for "Efficient in vivo genome editing mediated by stem cells-derived extracellular vesicles carrying designer nucleases"

Bobis-Wozowicz et al.

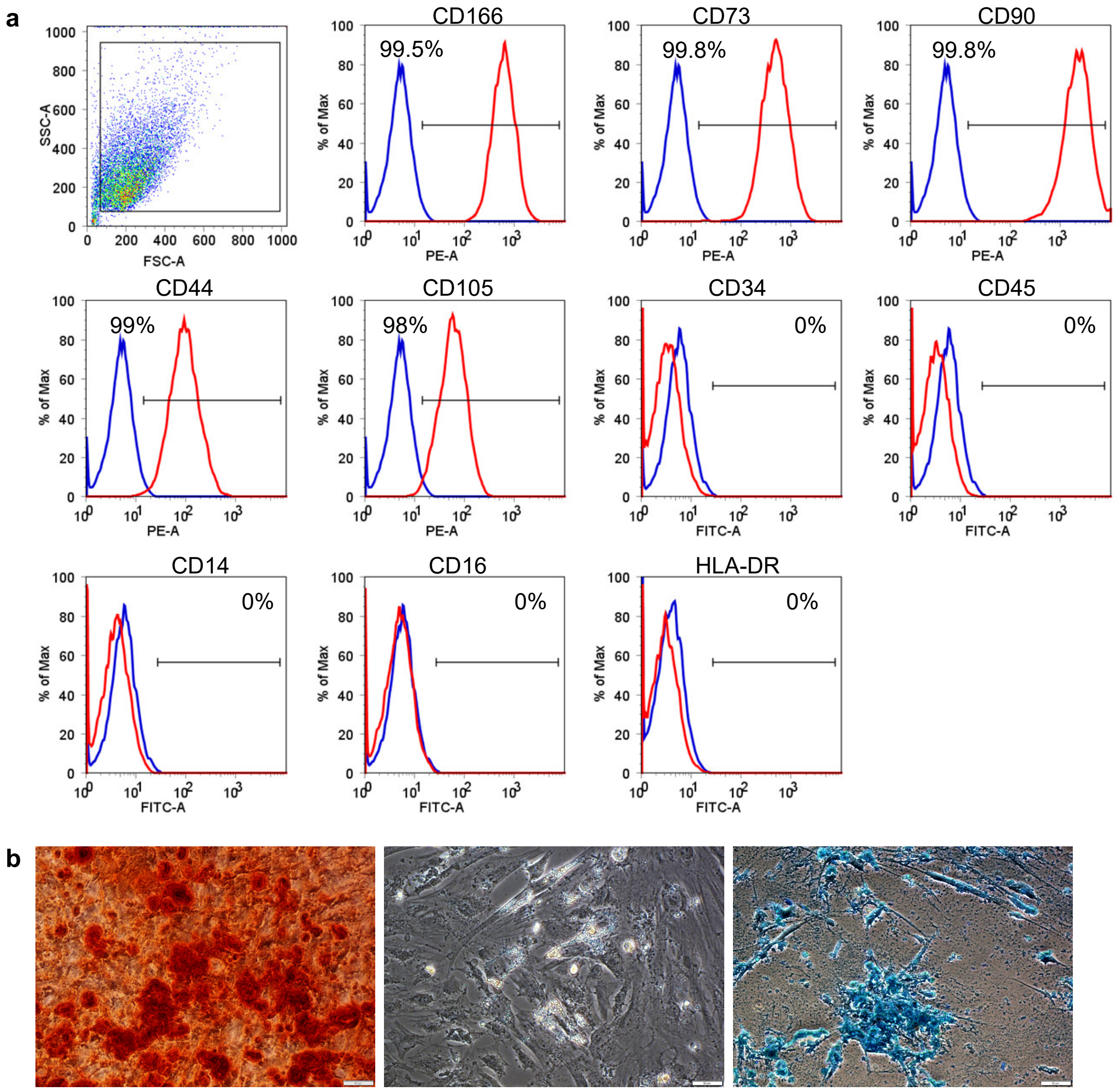


**Supplementary Figure 1.** Characterization of UC-MSCs. **a.** Expression of surface markers on UC-MSCs analyzed by flow cytometry. The percentage of cells expressing antigens typical for MSC (CD166, CD73, CD90, CD44, CD105) and non-hematopoietic nature (CD34, CD45, CD14, CD16, HLA-DR) was measured. Representative histograms are shown. Blue line – isotype control, red line – cells stained with respective antibody. **b.** UC-MSC differentiation into osteocytes (left), adipocytes (middle) and chondrocytes (right). Representative pictures are shown. Scale bar is 50 μm.

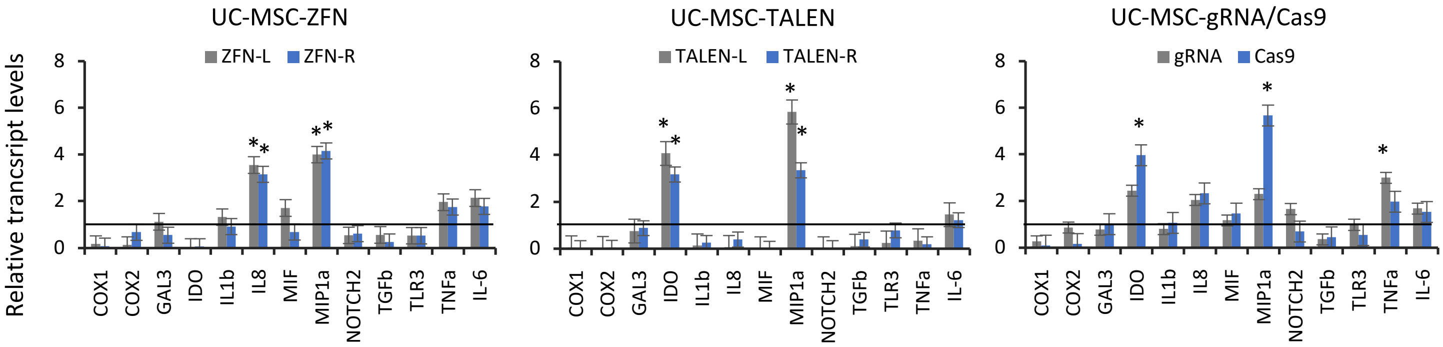


**Supplementary Figure 2. Analysis of expression levels of genes related to immunomodulation in UC-MSC stably expressing designer nucleases.** Series of genes related to inflammation (*COX1, COX2, IL-1b, IL-8, MIP1a, TNFa*), immunomodulation (*Gal-3, NOTCH2, TLR3*), immunosupression (*IDO*) and anti-inflammatory status (*MIF, IL-6*) were analyzed using real time qPCR method. Gene expression changes in genetically modified UC-MSC cell lines expressing ZFN-L/R (left), TALEN-L/R (middle) or gRNA/Cas9 (right) relative to unmodified cells (WT) are shown. Results are presented as the mean ± SD from two expreriments run in duplicate. Statistical significance at $p \leq 0.05$ is indicated with an asterisk.

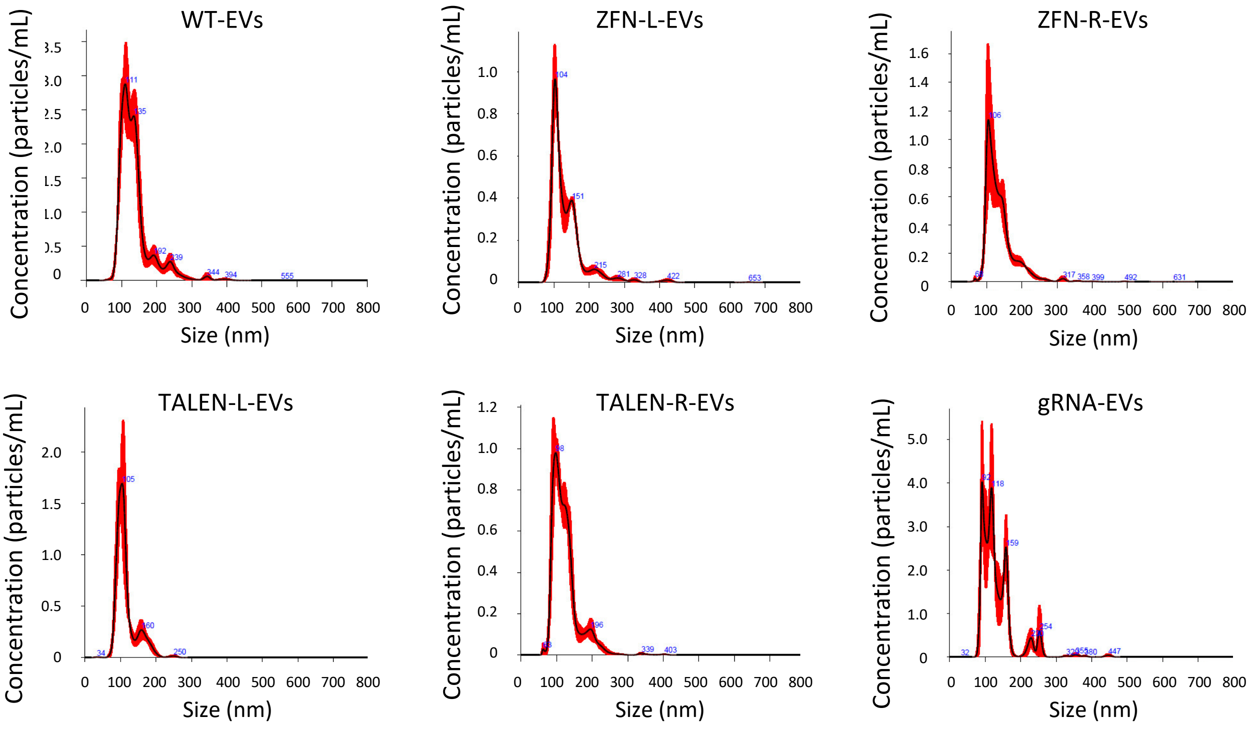


**Supplementary Figure 3.** Nanoparticle Tracking Analysis (NTA) of UC-MSC-DN-EVs. Representative histograms for each EV type are shown.

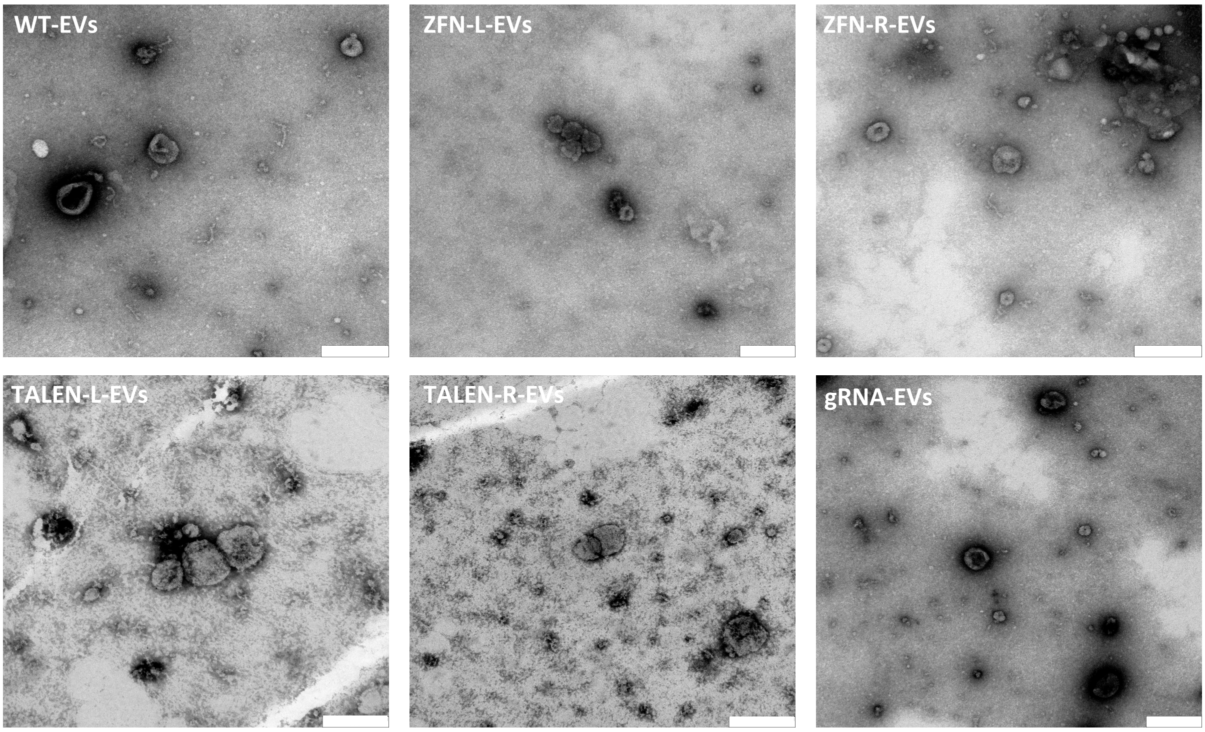


**Supplementary Figure 4.** Visualization of UC-MSC-DN-EVs using Transmission Electron Microscopy (TEM). Representative pictures for each EV type are shown. Scale bar is 200 nm.

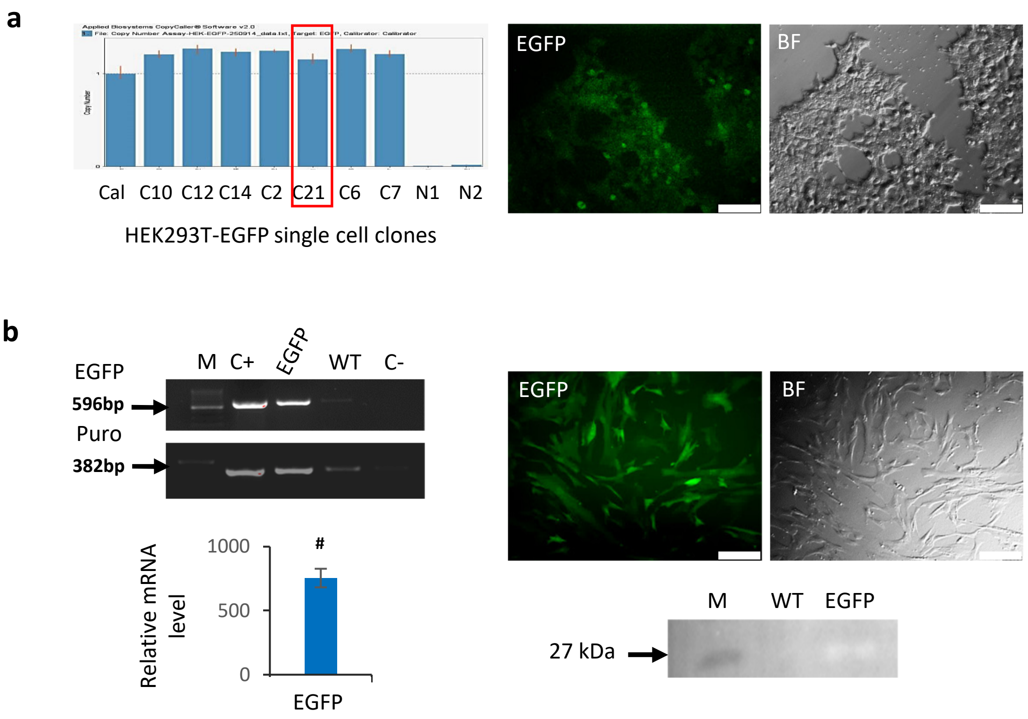


**Supplementary Figure 5.** Generation of EGFP-expressing cell lines. Two cell types: HEK293T and UC-MSCs were genetically modified by lentiviral transduction with EGIP vector to stably express EGFP gene. **a.** Selection of HEK293T-EGFP cell line containg a single copy of EGFP gene. Left panel: 7 single cell-derived clones were analyzed by CopyCaller Software v2.0 (Applied Biosystems) and the cell clone C21 was selected for further studies (indicated by a red frame). Right panel: microscopic pictures of HEK293T-EGFP C21 clone in green fluorescence and in bright field. Scale bar is 100 µm. **b**. Analysis of UC-MSC-$EGFP^{dim}$ polyclonal cell line. Left panel: PCR-based genotyping to confirm integration of EGFP gene and puromycin (puro) resistance gene (upper part) and measurement of mRNA expression level by real time qPCR method, relative to unmodified cells (lower part). Statistical significance at $p<0.001$ is indicated with $^{\#}$. Right panel: representative pictures of UC-MSC-EGFP cells in fluorescence microscopy and in bright field. Scale bar is 100 µm (upper part). Detection of EGFP protein in UC-MSC-EGFP cells by Western blot (lower part). Abbreviations: M – marker; WT – wild type; EGFP – EGFP-expressing cell line.

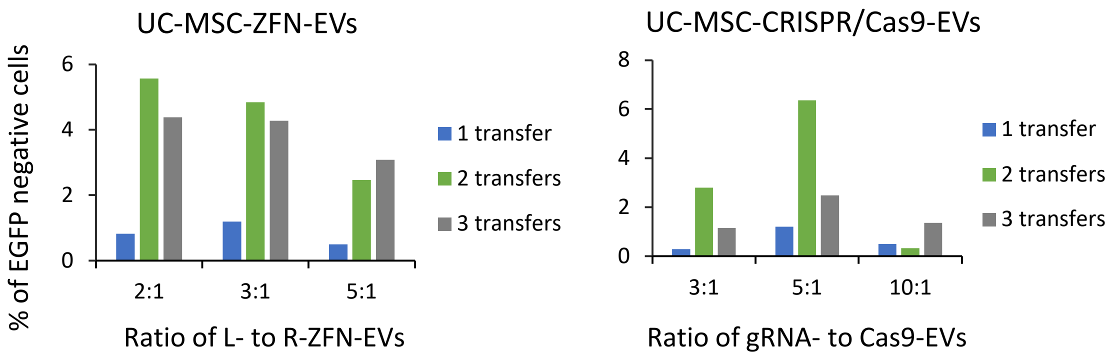


**Supplementary Figure 6.** Establishing the ratio of designer nucleases domains for efficient genome editing. HEK293T-EGFP cells were treated up to three times with EVs containing either ZFN-L and ZFN-R or gRNA and Cas9 nucleases at different ratios. After 5 days upon the last EV transfer the cells were collected and EGFP knockout was measured by flow cytometry.

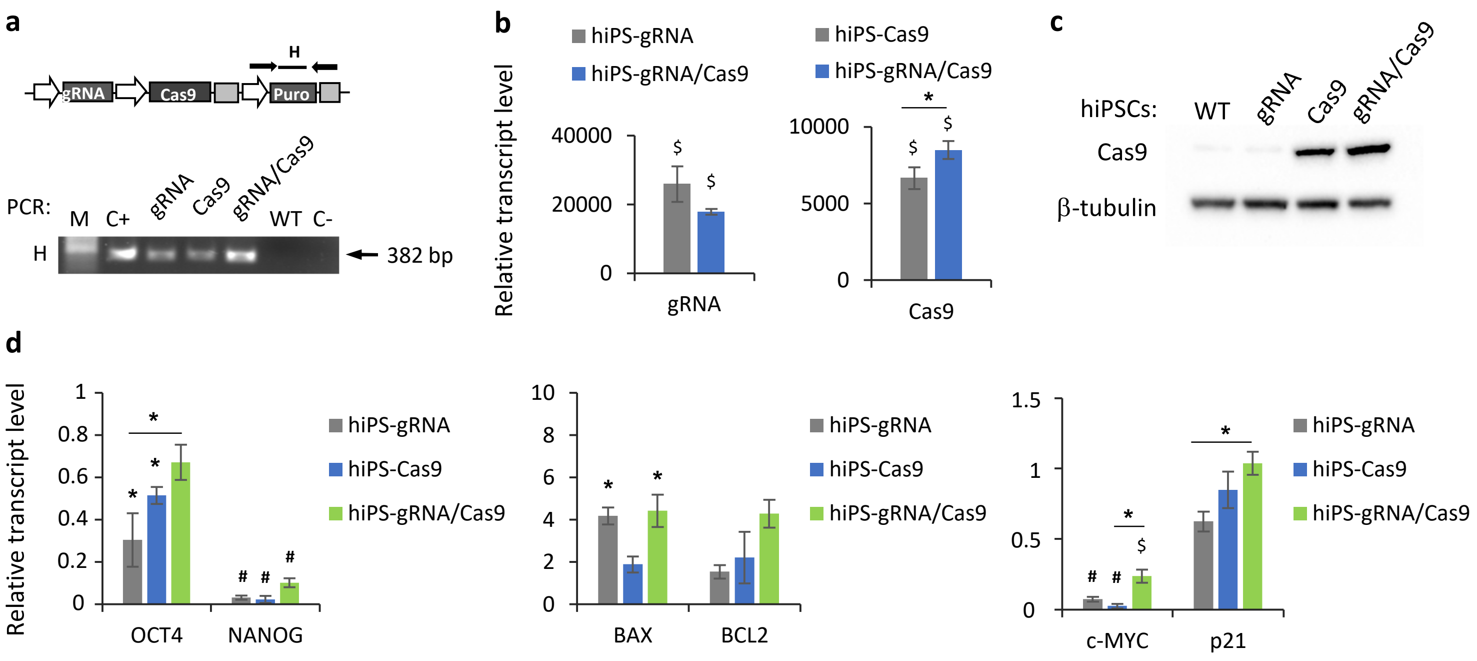


**Supplementary Figure 7.** Characteristics of isogenic hiPSC lines expressing different components of the CRISPR/Cas9 system targeting the EGFP gene. **a.** Analysis of stable integration of puromycin restance gene in hiPSC lines. Upper part: schematic representation of the expression cassettes. Lower part: agarose-gel electrophoresis of the amplicons obtained in PCR-based genotyping. **b.** Transcript levels for gRNA and Cas9 in different hiPSC lines measured with real time qPCR. The data were analysed using the ΔΔCt method and β2-microglobulin as endogenous control. Statistical significance at $p<0.05$ and $p<0.01$ is indicated with * or $^{\$}$, respectively. **c.** Detection of Cas9 protein by Western blot. Equal loading was confirmed by detection of β-tubulin. **d.** Analysis of mRNA levels for genes related to pluripotency (OCT4, NANOG), apoptosis (BAX, BCL-2), and proliferation status (c-MYC, p21) in genetically modified hiPSC lines in comparison to WT cells. Analysis was done with real time qPCR method using the ΔΔCt method and β2-microglobulin as endogenous control. Results show the mean ± SD from two experiments performed in duplicate. Statistical significance at $p<0.05$, $p<0.01$ and $p<0.001$ is indicated with *, $^{\$}$ or $^{\#}$, respectively.

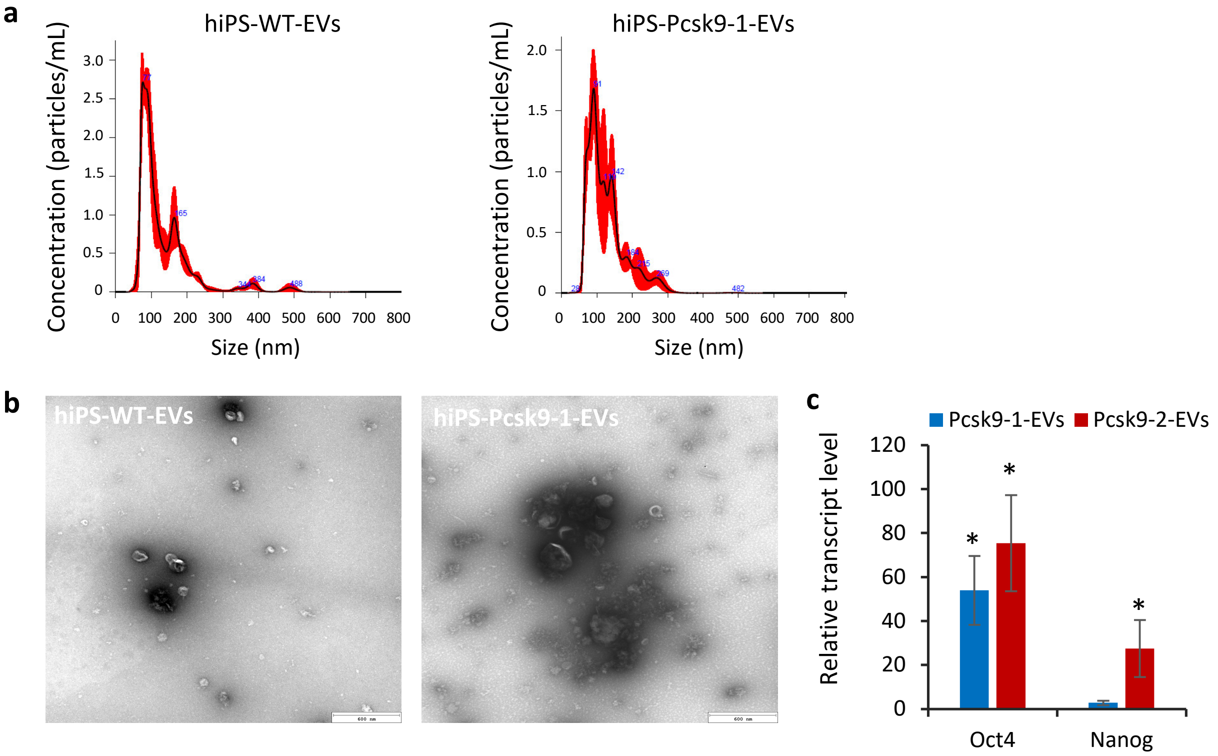


**Supplementary Figure 8.** Characteriazation of hiPS-EVs carrying the CRISPR/Cas9 system targeting mouse Pcsk9 gene. a. Represeatative histograms of EVs size distribution and concentration measured by NanoSight. b. Representative images of EV by transmission electron microscopy. Scale bar shows 600 nm.

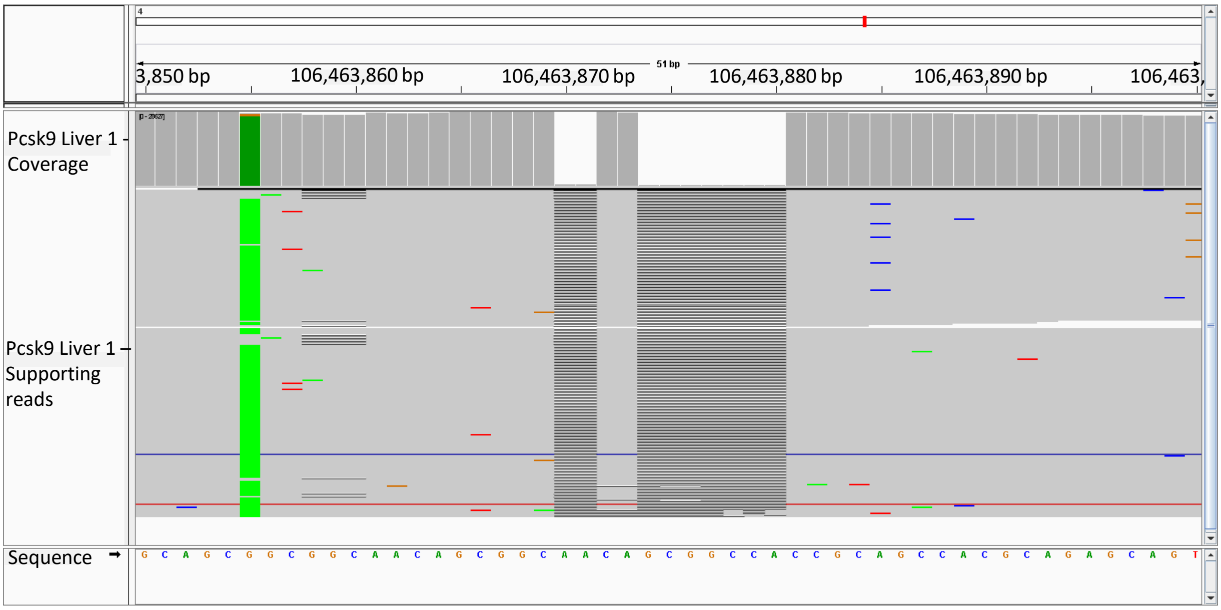


**Supplementary Figure 9.** Identification of indels in Pcsk9 gene in mouse liver upon hiPS-CRISPR/Cas9-EVs injection by next generation sequencing (NGS). Representative image showing a part of the alignment of the sequences obtained by Illumina sequencing to the mouse genome. Central part of the picture shows a 2 nt deletion followed by 7 nt deletion at the positions 106,463,869 and 106,463,871, respectively.

**Supplementary Table 1**. Primers used for genotyping of UC-MSC-DNs and iPS-DNs.

| PCR | Detected fragment | Primer forward (5' – 3') | Primer reverse (5' – 3') | Amplicon length (bp) |
|---|---|---|---|---|
| A | Junction of EF1a promoter and ZFN | TTTGCCCTTTTTGAGTTTGG | GATAGGCAGGTTGTAGCCGC | 788 |
| B | Junction of ZFN and TKpA | GAACTGATCGAAATCGCCCG | TGGGCCTTCACCCGAACTT | 749 |
| C | Junction of TALEN and TKpA | ATCAAGGAAACCGGACGGAG | TGGGCCTTCACCCGAACTT | 672 |
| D | PGK –puro-pA cassette | GGGGTTGGGGTTGCGCCT | GCCTGCTATTGTCTTCCCAA | 1609 |
| E | Junction of U6 promoter and gRNA | ATCGATGCTAGCTGTACAAAAAAGC | GACTCGGTGCCACTTTTTCA | 421 |
| F | Junction of EF1a promoter and SpCas9 | TTTGCCCTTTTTGAGTTTGG | GTGATTTGGCGGGTTTCCAC | 2972 |
| G | Junction of SpCas9 and TKpA | AAAGTGCTTACACGCTCGGA | TGGGCCTTCACCCGAACTT | 1931 |
| H | Puromycin resistance gene | GTCACCGAGCTGCAAGAACT | GCTCGTAGAAGGGGAGGTTG | 382 |
| I | SpCas9 | TCAGACCACACAGAAAGGGC | TTCGGATTTCGCCATTTGCC | 875 |

**Supplementary Table 2**. Size and concentration of UC-MSC-DN-EVs measured with NanoSight. SE – standard error.

| EV type | Mean ± SE [nm] | Mode ± SE [nm] | EV yield from 90 ml CM ± SE [x $10^{10}$] |
|---|---|---|---|
| WT | 135.2 ± 1.8 | 109.5 ± 4.8 | 5.8 ± 0.1 |
| ZFN-L | 135.9 ± 1.9 | 109.9 ± 1.2 | 6.2 ± 0.36 |
| ZFN-R | 132.4 ± 3.4 | 110.25 ± 8.3 | 10 ± 1.2 |
| TALEN-L | 125.6 ± 8.7 | 99.5 ± 4 | 12.6 ± 0.5 |
| TALEN-R | 130.3 ± 8.7 | 105.4 ± 5.3 | 9.4 ± 0.5 |
| gRNA | 133.3 ± 2.2 | 103.35 ± 9 | 6.75 ± 0.3 |
| Cas9 | 127.9 ± 4.7 | 103.5 ± 6.1 | 6.8 ± 0.78 |

**Supplementary Table 3.** Indels identified in EGFP gene by NGS in mouse liver samples after UC-MSC-DN-EVs treatment. Reads with minimal frequency of 10% were considered. DP – raw read depth; SR – number of supporting reads.

| Sample | EV type | Indel position | Indel type | Indel length [nt] | DP | SR | % of reads |
|---|---|---|---|---|---|---|---|
| Liver-1 | ZFN-EV | 514 | ins | 4 | 24729 | 6956 | 28.1 |
| Liver-2 | ZFN-EV | 514 | ins | 4 | 30254 | 6904 | 22.8 |
| Liver-3 | ZFN-EV | 514 | ins | 4 | 31614 | 14520 | 45.9 |
| Liver-4 | ZFN-EV | 514 | ins | 4 | 41549 | 19433 | 46.8 |
| Liver-5 | TALEN-EV | 207 | del | 3 | 79688 | 33592 | 42.1 |
| | TALEN-EV | 222 | del | 1 | 80804 | 34124 | 42.2 |
| | TALEN-EV | 227 | ins | 1 | 80968 | 34122 | 42.1 |
| Liver-6 | TALEN-EV | 207 | del | 6 | 47526 | 9788 | 20.6 |
| | TALEN-EV | 214 | del | 1 | 47399 | 9612 | 20.3 |
| | TALEN-EV | 216 | del | 7 | 47427 | 9614 | 20.3 |
| Liver-7 | TALEN-EV | 207 | del | 5 | 25824 | 3367 | 13 |
| Liver-8 | CRISPR/Cas9-EV | 173 | del | 10 | 42018 | 20395 | 48.5 |
| Liver-9 | CRISPR/Cas9-EV | 173 | ins | 2 | 28508 | 12346 | 43.3 |
| Liver-10 | CRISPR/Cas9-EV | 170 | del | 6 | 46923 | 18643 | 39.7 |

**Supplementary Table 4**. Size and concentration of hiPS-Pcsk9-EVs measured with NanoSight. SE – standard error.

| **EV type** | **Mean ± SE [nm]** | **Mode ± SE [nm]** | **EV yield from 90 ml CM ± SE [x $10^{10}$]** |
|---|---|---|---|
| WT | 132.3 ± 2.8 | 79 ± 3.1 | 5.4 ± 0.05 |
| Pcsk9-1 | 128.2 ± 4 | 85.1 ± 9.4 | 6.1 ± 0.12 |
| Pcsk9-2 | 126.2 ± 7.6 | 101.3 ± 9.3 | 4.9 ± 0.06 |

**Supplementary Table 5**. Indels identified in mouse *Pcsk9* gene by NGS in mouse liver samples after hiPS-Pcsk9-2-EVs treatment. Reads with minimal frequency of 10% were considered. DP – raw read depth; SR – number of supporting reads.

| Sample | EV type | Indel position (Ch4) | Indel type | Indel length [nt] | DP | SR | % of reads |
|---|---|---|---|---|---|---|---|
| Liver-1 | Pcsk9-2-EV | 106463868 | del | 2 | 71172 | 34568 | 48.6 |
| | Pcsk9-2-EV | 106463872 | del | 7 | 70210 | 34527 | 49.1 |
| Liver-2 | Pcsk9-2-EV | 106463868 | del | 9 | 64897 | 31810 | 49 |
| Liver-3 | Pcsk9-2-EV | 106463868 | del | 2 | 51012 | 25055 | 49.1 |
| | Pcsk9-2-EV | 106463872 | del | 7 | 50096 | 24916 | 49.7 |
| Liver-4 | Pcsk9-2-EV | 106463868 | del | 9 | 55219 | 26754 | 48.5 |
| Liver-5 | Pcsk9-2-EV | 106463868 | del | 8 | 57142 | 28087 | 49.2 |
| | Pcsk9-2-EV | 106463872 | del | 1 | 56186 | 28024 | 49.9 |
| Liver-6 | Pcsk9-2-EV | 106463868 | del | 9 | 47495 | 22814 | 48 |
| Liver-7 | Pcsk9-2-EV | 106463868 | del | 8 | 61669 | 30266 | 49.1 |
| | Pcsk9-2-EV | 106463877 | del | 1 | 60796 | 30266 | 49.8 |
| Liver-8 | Pcsk9-2-EV | 106463868 | del | 9 | 61111 | 29489 | 48.3 |
| Liver-9 | Pcsk9-2-EV | 106463872 | del | 7 | 55448 | 27456 | 49.5 |

**Supplementary Table 6.** Oligonucleotide sequences for targeting mPrkdc gene.

| **Oligo name** | **Sequence (5' – 3')** | **Purpose** |
|---|---|---|
| GeneArt-mPcsk9-1-F | TAATACGACTCACTATA**G**CAGAGCAGTGGGTGC | gRNA synthesis |
| GeneArt-mPcsk9-1-R | TTCTAGCTCTAAAACATGGGCACCCACTGCTCT | gRNA synthesis |
| GeneArt-mPcsk9-2-F | TAATACGACTCACTATA**G**GGCGGCAACAGCGGC | gRNA synthesis |
| GeneArt-mPcsk9-2-R | TTCTAGCTCTAAAACCTGTTGCCGCTGTTGCCG | gRNA synthesis |
| mPcsk9-oligo1-F | atccGCAGAGCAGTGGGTGCCCAT | Cloning to LV vector |
| mPcsk9-oligo1-R | aaacATGGGCACCCACTGCTCTGC | Cloning to LV vector |
| mPcsk9-oligo2-F | atccGGCGGCAACAGCGGCAACAG | Cloning to LV vector |
| mPcsk9-oligo2-R | aaacCTGTTGCCGCTGTTGCCGCC | Cloning to LV vector |

**Supplementary Table 7.** Primer list used in RT-qPCR.

| Gene name | Primer forward [5'-3'] | Primer reverse [5'-3'] |
|---|---|---|
| hNANOG | ACCTCAGCTACAAACAGGTGAAG | TTCTGCGTCACACCATTGCT |
| hSOX2 | GGGAAAGTAGTTTGCTGCCTC | CAGGCGAAGAATAATTTGGGGG |
| hc-MYC | TCTCCGTCCTCGGATTCTCT | TTCTTGTTCCTCCTCAGAGTCG |
| hCOL1A1 | GGACACAGAGGTTTCAGTGGT | GCACCATCATTTCCACGAGC |
| hMSX1 | AAAGTGGCTGGAAGAGTCCC | GACACCGATTTCTCTGCGCT |
| hBAX | TTTTGCTTCAGGGTTTCATCCAG | CGGAAAAAGACCTCTCGGGG |
| hBCL2 | GATAACGGAGGCTGGGATGC | TGACTTCACTTGTGGCCCAG |
| ZFN | GAACTGATCGAAATCGCCCG | GATAGGCAGGTTGTAGCCGC |
| TALEN | ATCAAGGAAACCGGACGGAG | ATCGTTGCATTTCATCTGCTTGG |
| gRNA | AGAAATAGCAAGTTAAAATAAGGCT | CGACTCGGTGCCACTTTT |
| Cas9-1 | GGGGAACACAGACCGTCATT | TTCGGTTCTTGCGACGTGTA |
| hCOX1 | GAGTTTGTCAATGCCACCT | CAACTGCTTCTTCCCTTTG |
| hCOX2 | TCCCTTGGGTGTCAAAGGTAAA | TGGCCCTCGCTTATGATCTG |
| hGAL3 | CCAAAGAGGGAATGATGTTGCC | TGATTGTACTGCAACAAGTGAGC |
| hIDO | CGCTGTTGGAAATAGCTTC | CAGGACGTCAAAGCACTGAA |
| hIL-1β | AGACATCACCAAGCTTTTTTGCT | GCACGATGCACCTGTACGAT |
| hIL-8 | TTAGCACTCCTTGGCAAAACTG | CTGGCCGTGGCTCTCTTG |
| hMIF | CAAGGCCAACCGCGAGAAGA | GGATAGCACAGCCTGGATAG |
| hMIP1α | ACTTGCCGGGAGGTGTAGCT | CAACCAGTTCTCTGCATCACTTG |
| hNOTCH2 | TGGGCTACACTGGGAAAAAC | TAGGCACTGGGACTCTGCTT |
| hTGFβ | AAGGCGAAAGCCCTCAATTT | CAGCAACAATTCCTGGCGATA |
| hTLR3 | AGGATTGGGTCTGGGAACA | AAAAACACCCGCCTCAAAG |
| hTNFα | CCTCTGATGGCACCACCAG | TCTTCTCGAACCCCGAGTGA |
| hIL-6 | TTCGGCAAATGTAGCATG | AATAGTGTCCTAACGCTCATAC |
| EGFP | CTACGGCAAGCTGACCCTGAA | GCCGTCCTCCTTGAAGTCGAT |
| hOCT4 | CCTTCGCAAGCCCTCATTTCA | CCCACAGAACTCATACGGCG |
| hp21 | CTCAGGGTCGAAAACGGCGG | CAGGCTTCCTGTGGGCGGAT |
| hβ2microglobulin | AATGCGGCATCTTCAAACCT | TGACTTTGTCACAGCCCAAGATA |
| mPcsk9 | GCTTCTGCTCCAGAGGTCATC | TGTGAGGTCCCACTCTGTGA |
| mβ2microglobulin | CATACGCCTGCAGAGTTAAGCA | GATCACATGTCTCGATCCCAGTAG |
| Cas9-2 | GATAACGGCAGCATCCCTCA | GGGGATGCGGAAAGTCAGAA |

### Supplementary Methods

#### Flow cytometry analysis of surface markers on UC-MSC

UC-MSCs at passage 3 were harvested using TrypLe Select Enzyme (Gibco), and the cell suspension was centrifuged at 200×g for 5 min. Cells were counted and $10^5$ of cells were stained with a specific fluorescently-labeled antibody directed towards the following antigens: CD166, CD73, CD90, CD44, CD105, and HLA-DR (all PE-conjugated from Biolegend) or CD34, CD45, CD14, and CD16 (FITC-conjugated from BD Biosciences). Staining was performed in 100 µL of staining buffer, composed of PBS supplemented with 2% FBS, for 30 min at 4°C in the dark. Next, cells were washed in PBS, resuspended in 300µL of staining buffer and the percentage of fluorescently labeled cells was measured using the BD LSRFortessa flow cytometer (BD Biosciences). The obtained data were analyzed with the FlowJo software (FlowJo).

#### Differentiation of UC-MSCs into osteocytes, adipocytes and chondrocytes

UC-MSCs from passage 3 were used in differentiation studies. Briefly, $3x10^3$ cells were seeded on 12-well plates and 2 days later the differentiation was initiated by culturing the cells in specific media: StemPro® Osteogenesis Differentiation Kit, StemPro® Adipogenesis Differentiation Kit and StemPro® Chondrogenesis Differentiation Kit (all from Gibco). Cells were maintained in differentiation media for 21 days, with medium change every 3–4 days. Subsequently, the cells were fixed in 4% paraformaldehyde and stained with 2% Alizarin Red (Sigma-Aldrich) to detect calcium phosphate deposits released by osteocytes, or were stained with 1% Alcian Blue solution (Sigma-Aldrich) to detect proteoglycans produced by chondrocytes. Adipogenic differentiation was evaluated by the presence of fat droplets.
